# An activation pathway governs cell wall polymerization by a bacterial morphogenic machine

**DOI:** 10.1101/359208

**Authors:** Patricia D. A. Rohs, Jackson Buss, Sue Sim, Georgia Squyres, Veerasak Srisuknimit, Mandy Smith, Hongbaek Cho, Megan Sjodt, Andrew C. Kruse, Ethan Garner, Suzanne Walker, Daniel Kahne, Thomas G. Bernhardt

**Author notes:** These authors contributed equally (Jackson Buss, Sue Sim). To whom correspondence should be addressed. Thomas G. Bernhardt, Ph.D., Harvard Medical School, Department of Microbiology and Immunobiology Boston, Massachusetts 02115.

## Abstract

Cell elongation in rod-shaped bacteria is mediated by the Rod system, a conserved morphogenic complex that spatially controls cell wall (CW) assembly. In *Escherichia coli*, alterations in a CW synthase component of the system called PBP2 were identified that overcome other inactivating defects. Rod system activity was stimulated in the suppressors in vivo, and purified synthase complexes with these changes showed more robust CW synthesis in vitro. Polymerization of the actin-like MreB component of the Rod system was also found to be enhanced in cells with the activated synthase. The results suggest an activation pathway governing Rod system function in which PBP2 conformation plays a central role in stimulating both CW glycan polymerization by its partner RodA and the formation of cytoskeletal filaments of MreB to orient CW assembly. An analogous activation pathway involving similar enzymatic components is likely responsible for controlling CW synthesis by the division machinery.

## INTRODUCTION

Bacterial cells typically surround themselves with a cell wall exoskeleton made of the heteropolymer peptidoglycan (PG). This structure is essential for cell integrity and understanding its biogenesis is of great practical significance because the pathway is a proven target for many of our most effective antibiotic therapies (Silver, 2013). The PG layer is also the major determinant of bacterial cell shape such that studies of PG assembly are also of fundamental importance for determining the mechanisms responsible for bacterial growth and morphogenesis (Typas, Banzhaf, Gross, & Vollmer, 2012).

PG is composed of long glycan strands with a disaccharide repeating unit of N-acetylmuramic acid (MurNAc)-β-1-4-N-acetylglucosamine (GlcNAc) and a pentapeptide stem attached to the MurNAc sugar (Höltje, 1998). The strands are polymerized by membrane-embedded PG glycosyltransferase (PGTase) enzymes using the lipid-linked disaccharide-pentapeptide precursor called lipid II. The polymerized glycans are then crosslinked via the formation of amide bonds between attached peptides by transpeptidase (TPase) enzymes. Several different types of synthases with these activities work together to build what ultimately becomes a cell-shaped polymer matrix that envelops the cytoplasmic membrane and protects it from osmotic lysis.

To direct PG matrix assembly during cell growth and division, rod-shaped bacteria employ two multi-protein synthetic machineries organized by cytoskeletal filaments (Typas et al., 2012). The Rod system (elongasome) utilizes the actin-like MreB protein to promote cell elongation and maintain cell shape, whereas the cytokinetic ring (divisome) uses the tubulin-like FtsZ protein to orchestrate cell division and the construction of the daughter cell poles. For many years, the main PG synthases of these machineries were thought to be the class A penicillin-binding proteins (aPBPs) (Typas et al., 2012). These bifunctional synthases possess both PGT and TP activity to make PG, and until recently, the PGT domain of aPBPs was the only known family of PG polymerases. This view of PG biogenesis was called into question by the discovery of PG polymerase activity for the SEDS (shape, elongation, division, and sporulation) family protein RodA of the Rod system (Meeske et al., 2016).

SEDS family proteins are widely distributed in bacteria (Henriques, Glaser, Piggot, & Moran, 1998; Meeske et al., 2016) and are known to form complexes with class B PBPs (bPBPs) (Fay, Meyer, & Dworkin, 2010; Fraipont et al., 2011), which are monofunctional TPases only thought to be capable of PG crosslinking. Thus, SEDS-bPBP complexes have been proposed to represent a second type of PGT/TP enzymatic system for PG synthesis, with FtsW-PBP3 and RodA-PBP2 functioning as the SEDS-bPBP pairs for the divisome and Rod system, respectively (Cho et al., 2016; Meeske et al., 2016). Although it remains possible that the SEDS-bPBP synthases work together with aPBPs in the same complexes, functional and localization studies suggest otherwise (Cho et al., 2016). In both *Escherichia coli* and *Bacillus subtilis*, the aPBPs have been shown to display distinct subcellular localization dynamics from Rod system components and to be dispensable for the activity of the machinery (Cho et al., 2016; Meeske et al., 2016). It has therefore been proposed that a RodA-PBP2 complex forms the core PG synthase of the Rod system, an idea supported by recent evolutionary co-variation analysis (Sjodt et al., 2018), and the finding that the aPBPs largely operate outside of the cytoskeletal system during cell elongation (Cho et al., 2016). A similar division of labor between aPBPs and FtsW-PBP3 may also be taking place during cytokinesis, but the relative contributions of the two types of synthases to the division process requires further definition.

The discovery that RodA is a PG polymerase raises many important questions about the function of the Rod system. Is the polymerase activity of this new synthase regulated, and if so, how is its activity controlled to maintain a uniform rod shape? Does RodA work with PBP2 as proposed, and and if so, how is the polymerase activity of RodA coordinated with the crosslinking activity of PBP2? Coupling of these activities is expected to be critical as it is disrupted by beta-lactam antibiotics as part of their lethal mechanism of action (Cho, Uehara, & Bernhardt, 2014). For example, the beta-lactam mecillinam blocks the TP activity of PBP2 while leaving the activity of RodA unaffected. As a result, RodA generates uncrosslinked glycan strands that are rapidly degraded, causing a futile cycle of PG synthesis and degradation that is cytotoxic (Cho et al., 2014; 2016). Thus, during its normal function, the Rod system is likely to possess a fail-safe that prevents RodA from initiating PG polymerization unless it is engaged with PBP2 to crosslink its product glycans. Finally, aside from MreB, RodA, and PBP2, the Rod system includes the additional proteins MreC, MreD, and RodZ. Despite their broad conservation throughout cell wall producing bacteria, even in non-rod-shaped organisms lacking MreB (Alyahya et al., 2009), the function of these additional Rod system components remains unclear.

In this report, we describe the discovery of PBP2 variants that suppress the growth and shape defects of *mreC* hypomorphs. One of the altered PBP2 variants was shown to hyperactivate cell wall synthesis by the Rod system in vivo and to stimulate the polymerase activity of RodA-PBP2 complexes in vitro. Furthermore, studies of Rod system localization dynamics in the mutant cells indicate that the PBP2 variant promotes the formation of active Rod complexes by enhancing MreB filament formation. Overall, our results define an activation pathway for the cell elongation machinery in which PBP2 plays a central role in both stimulating PG polymerization by RodA and modulating MreB polymerization to orient new synthesis (Hussain et al., 2018). This mode of activation provides a built-in mechanism for coupling cell wall polymerization and crosslinking to prevent the toxic accumulation of uncrosslinked glycans. Moreover, the phenotypes of previously described cell division mutants (Du, Pichoff, & Lutkenhaus, 2016; Modell, Hopkins, & Laub, 2011; Modell, Kambara, Perchuk, & Laub, 2014) and our recent biochemical studies of FtsW in a complex with its cognate bPBP (Taguchi et al., 2018) suggest that this activation pathway is conserved to control PG synthesis by the divisome.

## RESULTS

### A strategy to identify mutants with hyperactive Rod systems

In *E. coli* and other organisms, each protein within the Rod system is required for proper functioning of the complex (Alyahya et al., 2009; Bendezú & de Boer, 2008; Bendezú, Hale, Bernhardt, & de Boer, 2009; Kruse, Bork-Jensen, & Gerdes, 2005; Leaver & Errington, 2005; Shiomi, Sakai, & Niki, 2008). Rod system defects result in a loss of rod shape and cell death under typical growth conditions, but spherical *E. coli* Rod^-^ mutants can survive on minimal medium at low temperatures (Bendezú & de Boer, 2008). Thus, mutants inactivated for the Rod system can be constructed under permissive conditions (minimal medium) and suppressors of these defects can be isolated by plating the mutants on rich medium (non-permissive conditions) and selecting for growth. Starting with a *ΔrodZ* mutant background, this suppressor isolation strategy has been successfully used to investigate how the interaction between RodZ and MreB may modulate Rod system function (Morgenstein et al., 2015; Shiomi et al., 2013). We reasoned that similar selections for suppressors of other Rod system defects might help us understand how the PG synthetic enzymes within the complex are controlled.

Defects resulting from from a missense mutation are expected to be easier for cells to overcome in a suppressor selection than those due to a deletion allele. We therefore developed a strategy to rapidly identify missense alleles in Rod system genes that result in a stable yet defective gene product. In a report that will be published separately, we applied this method to *mreC*. Several defective *mreC* alleles were identified, with the two mutants displaying the most severe defects encoding MreC proteins with a G156D or an R292H substitution (Figure 1A). When the *mreC(G156D)* or *mreC(R292H)* alleles were constructed at the native *mre* locus, the resulting cells displayed a morphological defect reminiscent of an *mreC* deletion (Figure 1B). Although stable MreC protein accumulated in these mutants (Figure 1C), the proteins were incapable of promoting Rod system activity. We therefore concluded that the MreC variants identified were functionally defective and therefore suitable for use in a suppressor analysis.

**Figure 1.**
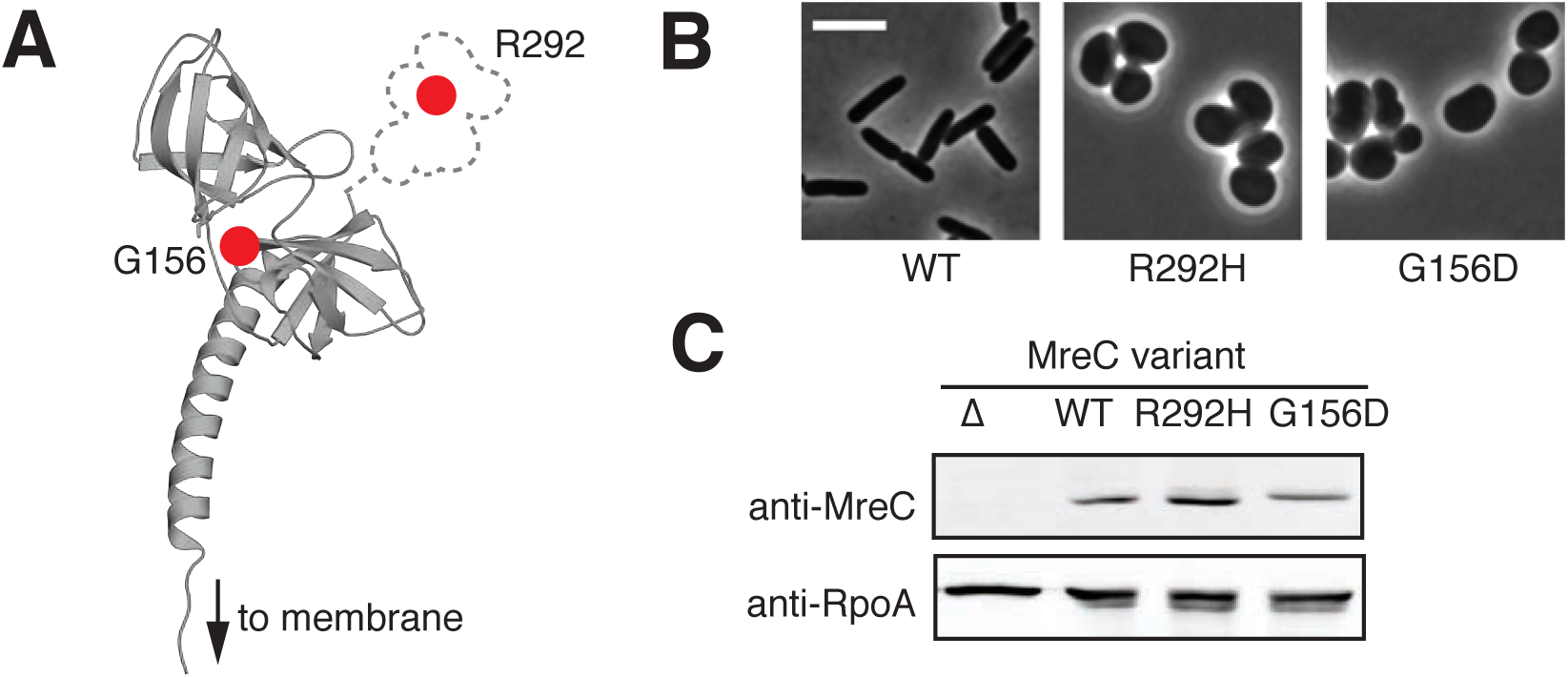
Amino acid substitutions in MreC affect protein function but not stability. **A.** *E. coli* MreC, modeled based on a crystal structure from *L. monocytogenes* using Phyre2 (Kelley & Sternberg, 2009; van den Ent et al., 2006). Locations of the amino acid substitutions affecting function are indicated by the red dots. **B.** Strains containing the indicated *mreC* point mutations at the native genomic locus [HC555, PR5, PR30] were grown overnight in M9 medium supplemented with 0.2% casamino acids and 0.2% glucose (M9 CAA glu), diluted to OD_600_=0.05 in the same medium, and grown at 30℃ until the OD_600_ reached 0.20. Cells were then gently pelleted and resuspended in LB to an OD_600_=0.025. Cells were then grown at 30℃ to an OD_600_ of 0.20. At this time, cells were fixed and imaged using phase-contrast microscopy. Scale bar, 5 μm. **C.** Immunoblot detecting MreC and the loading control RpoA. Each lane was loaded with 5 μg of total protein from whole cell extracts of *ΔmreC* [MT4], WT [HC555], *mreC(R292H)* [PR5], and *mreC(G156D)* [PR30] strains harvested in exponential-phase (OD_600_ ∼ 0.3).

Cells harboring the *mreC(G156D)* or *mreC(R292H)* alleles were plated on rich medium, the non-permissive condition for mutants defective for Rod system activity. Suppressors restoring growth arose at a frequency of 10^−5^. The majority of these isolates remained spherical, indicating that they had likely acquired mutations that allow spheres to grow on rich medium. However, visual screening identified several isolates that grew with a long axis, indicating at least a partial restoration of rod shape. Of these suppressors, two displayed near normal rod shape and were chosen for further analysis.

### Amino acid substitutions in PBP2 suppress MreC defects

Whole-genome sequencing was used to map the location of the *mreC* suppressor mutations. Both isolates harbored mutations in the *pbpA* (*mrdA*) gene encoding PBP2, the PG crosslinking enzyme of the Rod system. Although the *pbpA(T52A)* allele was originally found to suppress *mreC(G156D)* and the *pbpA(L61R)* allele was first isolated as a suppressor of *mreC(R292H),* neither suppressor was allele specific. Both were capable of suppressing the shape and viability defects of either *mreC* allele when the mutants were reconstructed in an otherwise normal parental strain background (Figure 2A-B). However, *pbpA(L61R)* was more robust at restoring normal rod shape than the *pbpA(T52A)* allele.

**Figure 2.**
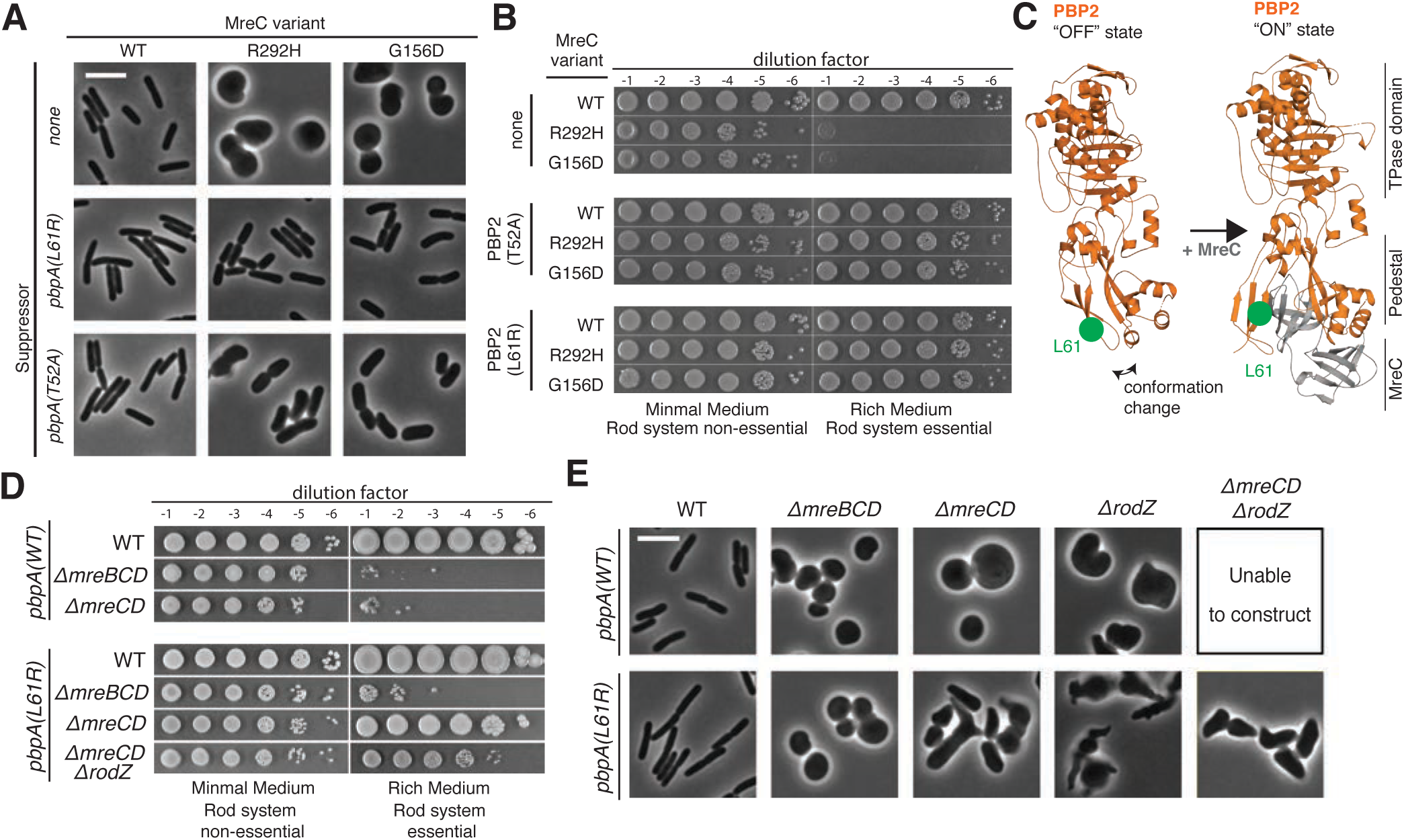
Substitutions in PBP2 suppress the growth and shape phenotypes of rod system mutants. **A.** Strains containing the indicated point mutations at the native genomic locus [PR164, PR165, PR166, PR127, PR128, PR129, PR131, PR124, PR125, PR161, PR162, PR163] were grown overnight in M9 supplemented with 0.2% casamino acids and 0.2% glucose (M9 CAA glu), diluted to OD_600_=0.05 in the same medium, and grown at 30℃ until the OD_600_ reached 0.20. Cells were then gently pelleted, then resuspended and diluted in LB such that the OD_600_ =0.025. Cells were allowed to grow at 30℃ until the OD_600_ reached 0.20. At this time, cells were fixed and imaged using phase-contrast microscopy. Scale bar, 5 μm. **B.** Overnight cultures of the above strains were serially diluted and spotted on either M9 CAA glu agar (Rod non-essential) or LB agar (Rod essential). Plates were incubated at 30℃ for either 40 h (M9) or 16 h (LB). **C.** Shown are E. coli PBP2 and the PBP2-MreC structures modeled from PDB-5LP4 and PDB-5LP5 (Contreras-Martel et al., 2017) using Phyre 2 (Kelley & Sternberg, 2009). PBP2 is orange with residue L61 in green. MreC is gray. **D.** Strains containing the indicated mutations were grown and spotted as in **B.** [Top to bottom: PR132, PR136, PR137, PR78, PR129, PR140, PR149]. **E.** The above strains were grown and prepared for phase-contrast microscopy as described in (A). Scale bar, 5 μm.

The changes in the altered PBP2 derivatives map to the membrane proximal region of the protein often referred to as the pedestal or non-penicillin-binding domain (Figure 2C). In the solved structures of bPBPs (Contreras-Martel, Dahout-Gonzalez, Martins, Kotnik, & Dessen, 2009; Han et al., 2010; Powell, Tomberg, Deacon, Nicholas, & Davies, 2009), this region consists of two interacting subdomains connected by a third subdomain forming a hinge that sits just underneath the catalytic TP domain. In a recently solved structure of an MreC-PBP2 complex from *Helicobacter pylori*, MreC interacts with the pedestal domain of PBP2 and in doing so causes its two interacting subdomains to swing open (Contreras-Martel et al., 2017) (Figure 2C). The alterations in PBP2 that suppress the MreC defects are not predicted to be at locations directly involved in the PBP2-MreC interface. Moreover, PBP2 derivatives with changes in the same region, PBP2(Q51L) and PBP2(T52N), were previously shown to suppress a Rod system defect caused by a Δ*rodZ* mutation (Shiomi et al., 2013). We therefore hypothesized that the conformational change in PBP2 induced by MreC may be part of a mechanism controlling PG synthesis by the core enzymatic components of the Rod system. We further reasoned that the PBP2 variants we identified might spontaneously achieve an activated conformation such that they bypass the normal requirement for MreC and other components of the Rod machinery that may have regulatory functions.

To begin testing our hypothesis, we assessed whether the strongest suppressor of *mreC* missense mutations, PBP2(L61R), could also suppress the shape and viability defects of mutants deleted for Rod system genes. This variant suppressed the growth defect of Δ*rodZ* cells and partially restored their shape as expected based on similarity to previously isolated Δ*rodZ* suppressors (Shiomi et al., 2013) (Figure 2D, E). PBP2(L61R) also had the additional ability to suppress the growth defect of a Δ*mreCD* mutant and a Δ*mreCD* Δ*rodZ* triple mutant (Figure 2D, E). Although rod shape was not fully restored in these cells, they displayed a long axis indicative of at least partial restoration of Rod system function (Figure 2E). Notably, this PBP2 variant was incapable of suppressing the shape or viability defects of a Δ*mreBCD* mutation (Figure 2D), indicating that the actin-like MreB protein remains essential for Rod system function in cells producing this altered PBP. These results are consistent with PBP2(L61R) adopting an activated conformation that mimics that induced upon assembly of the complete Rod system. Furthermore, the observation that partial rod shape can be restored with just MreB, RodA, and a PBP2 variant suggests that these three proteins form the minimal and essential core of the system.

### PBP2(L61R) activates cell wall synthesis by the Rod system

The hypothesis that PBP2(L61R) is an activated variant of PBP2 predicts that cells harboring the altered protein should have elevated Rod system activity. To investigate this possibility, the *pbpA(L61R)* allele was engineered into cells with an otherwise normal complement of Rod system components. The growth rate of these cells was indistinguishable from that of wild type in both rich and minimal medium (Table 1). However, the PBP2(L61R) cells were ∼20% longer and ∼10% thinner than cells with PBP2(WT) (Table 1), providing an early indication that the Rod system may be activated by the altered PBP2. To monitor Rod system activity more directly, we followed cell wall synthesis in cells radiolabeled with [^3^H]-*meso*-diaminopimelic acid (mDAP), an amino acid unique to the PG stem peptide. For these studies, we used a previously described genetic background in which the divisome can be inactivated by an inducible copy of the FtsZ antagonist SulA and aPBP activity can be inhibited by the thiol-reactive reagent (2-sulfanatoethyl)methanethiosulfonate (MTSES) (Cho et al., 2016). Thus, when SulA is produced and MTSES is added, radiolabel incorporation is mediated principally by the Rod system and thus reflects its activity (Figure 3A).

**Figure 3.**
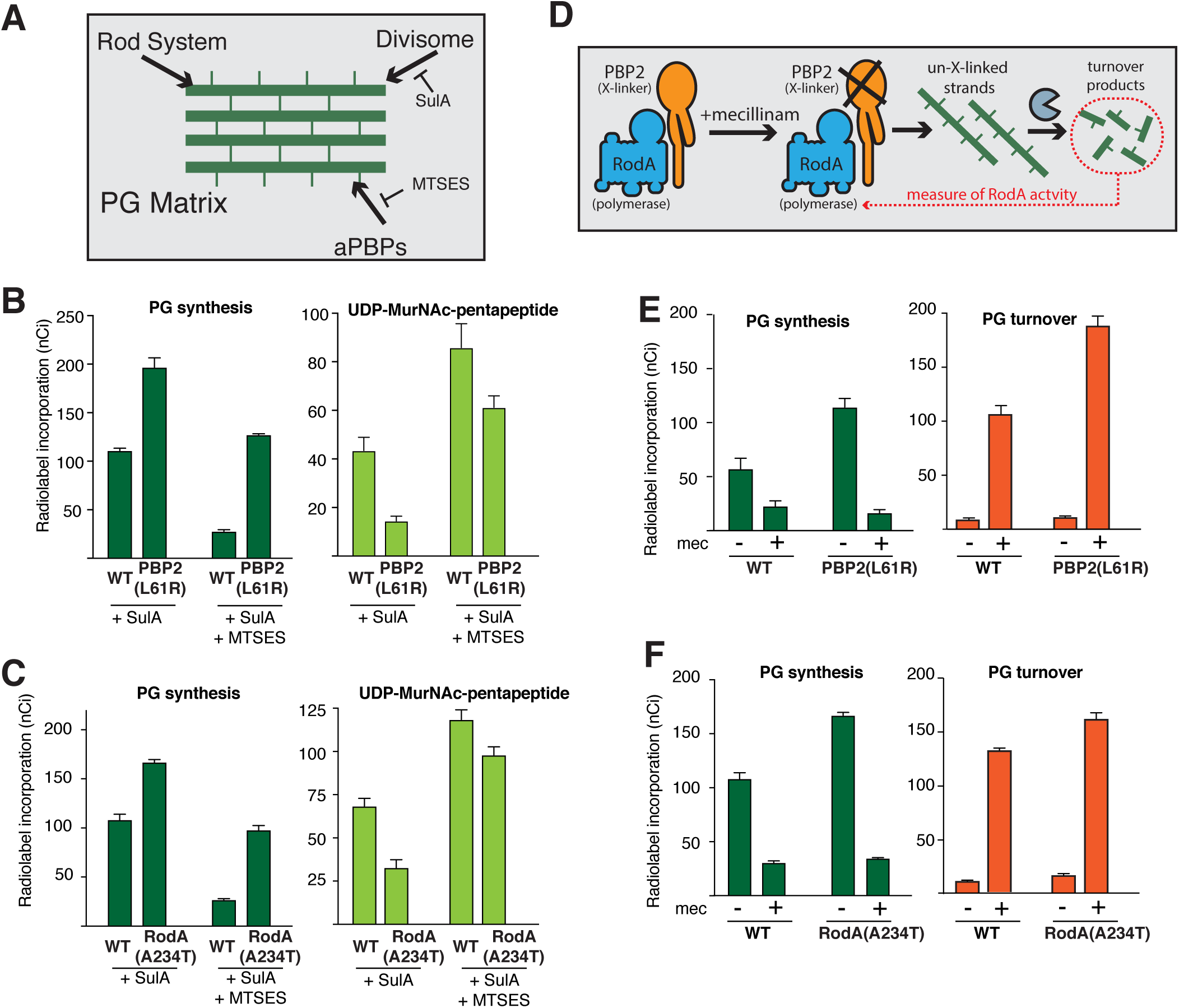
Cells expressing PBP2(L61R) or RodA(A234T) synthesize more peptidoglycan than wild type. **A.** Schematic of treatments used to inhibit specific PG synthesis systems during labeling experiments. In the labeling strains all aPBPs have been deleted (Δ*mtgA* Δ*mrcA* Δ*pbpC*) except for PBP1b, in which a cysteine mutation residue has been engineered near the active site (*mrcB(S247C)*), rendering it sensitive to MTSES. Labeling strains also contain mutations to block peptidoglycan recycling (Δ*lysA,* Δ*ampD*), and an attached plasmid to express the FtsZ inhibitor SulA under inducible control (attHKHC859). **B.** Labeling strains encoding PBP2(WT) or PBP2(L61R) at the native genomic locus [PR116(attHKHC859) and PR117(attHKHC859)] were pre-treated with 1.5 mM IPTG to induce SulA production and 1 mM MTSES, as indicated. Strains were then pulse-labelled with [^3^H]-mDAP, and peptidoglycan precursors (UDP-MurNAC-pentapeptide) and synthesis were measured. Results are the average of three independent experiments. Error bars represent the standard error of the mean. **C.** The same experiments and analysis as in (B) were performed using labeling strains encoding RodA(WT) or RodA(A234T) at the native genomic locus [PR146(attHKHC859) and PR147(attHKHC859)]. **D.** Schematic illustrating the mechanism of turnover-product production upon beta-lactam treatment. **E.** Labeling strains encoding PBP2(WT) or PBP2(L61R) at the native genomic locus [PR116(attHKHC859) and PR117(attHKHC859)] were pre-treated with 1.5 mM IPTG to induce SulA production. The indicated samples were also pre-treated with 10 μg/mL mecillinam. Strains were then pulse-labelled with [^3^H]-mDAP, and peptidoglycan synthesis and turnover products (anhydroMurNAC-tripeptide and -pentapeptide) were measured. Results are the average of four independent experiments. Note that a different stock of [^3^H]-mDAP was used for these experiments than in other panels such that total labeling observed was lower. **F.** The same experiments and analysis as in (E) were performed using labeling strains encoding RodA(WT) or RodA(A234T) at the native genomic locus [PR146(attHKHC859) and PR147(attHKHC859)]. Results are the average of three independent experiments.

**Table 1.**
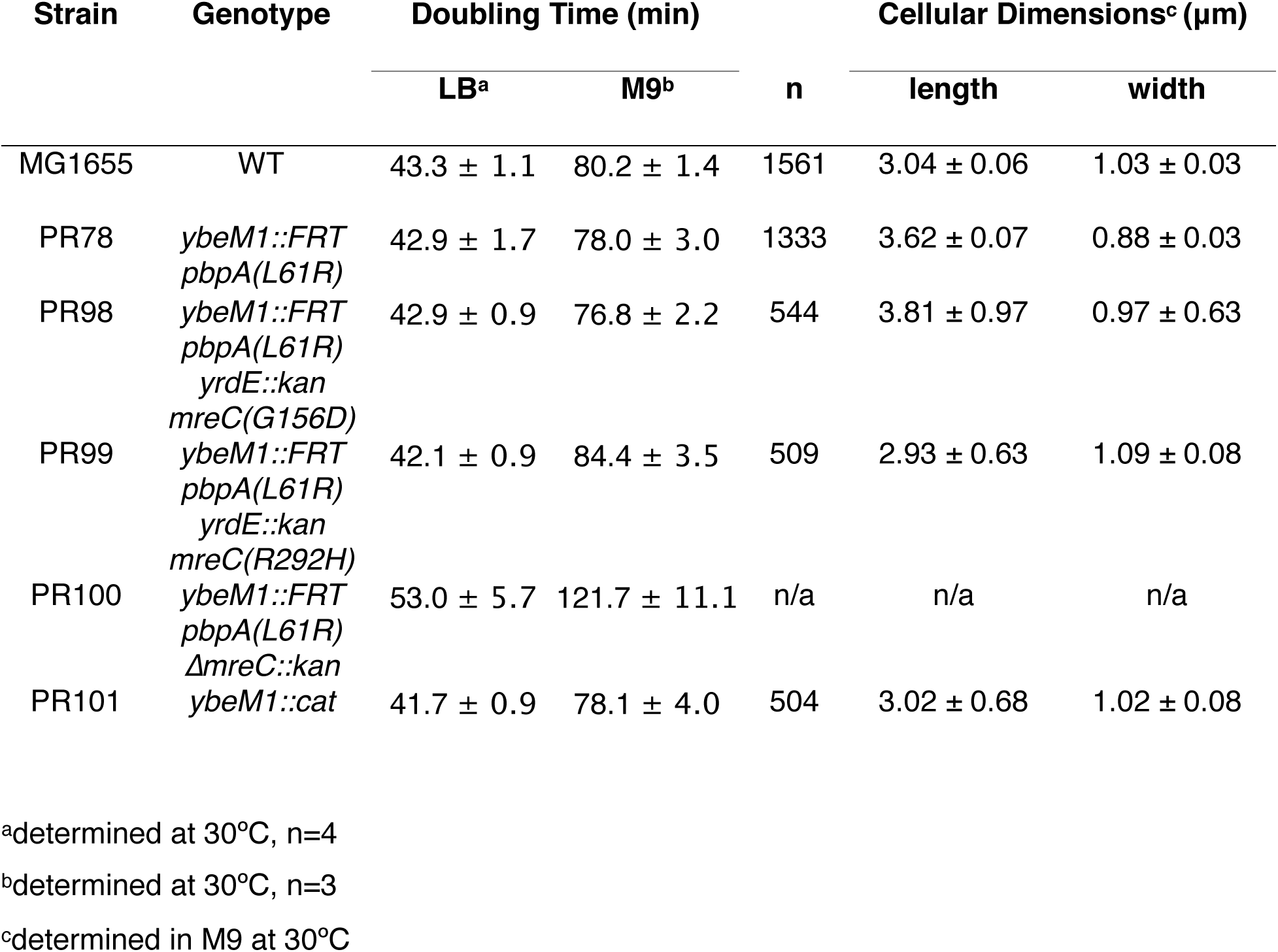
Growth rate and morphology of strains with altered Rod system proteins

Following divisome inhibition, PBP2(L61R) cells synthesized PG at approximately twice the rate of wild-type cells (197±10 nCi vs. 111±2 nCi over ten minutes, p<0.0001, Figure 3B). This increased synthesis activity was retained upon MTSES inhibition of the aPBPs, indicating that it indeed reflected elevated PG incorporation by the Rod system (127±1 nCi vs. 37.1±0.3 nCi over 10 minutes, p<0.0001, Figure 3B). The increase radiolabel incorporation into PG was also accompanied by a corresponding decrease in the labeled pool of the precursor UDP-MurNAc-pentapeptide, indicating that flux through the PG synthesis pathway is likely increased in the PBP2(L61R) cells (Figure 3B). Immunoblot analysis and labeling with the fluorescent penicillin derivative Bocillin failed to detect any changes in MreB or PBP2 levels in cells harboring the altered PBP2 protein (Figure 3, **supplement 1**). We therefore conclude that PBP2(L61R) is most likely activating PG synthesis by stimulating the activity of the Rod system.

### Rod system activation involves the stimulation of PG polymerization by RodA

In addition to changes in PBP2, RodA variants RodA(A234T) and RodA(T249P) were also previously identified as suppressors of a Δ*rodZ* mutation (Shiomi et al., 2013). We reconstructed the *rodA*(A234T) mutant at its native locus and confirmed this suppression activity and that the change in RodA was also capable of suppressing the growth and shape defects of the MreC variants we isolated, MreC(G156D) and MreC(R292H) (Figure 4). However, RodA(A234T) could not compensate for the deletion of Rod system genes other than *rodZ*, indicating that it is not as potent of a suppressor as PBP2(L61R) (Figure 4). Nevertheless, the suppression results suggested that RodA(A234T) is also capable of activating PG synthesis by the Rod system. We therefore monitored PG synthesis in *rodA*(A234T) mutant cells and found that Rod system activity was indeed enhanced relative to wild-type (167±3 nCi vs. 108±6 over ten minutes, p=0.001, Figure 3C). In line with the relative suppression power of the variants, the observed PG synthesis activation by RodA(A234T) was not as great as that observed in cells producing PBP2(L61R).

**Figure 4.**
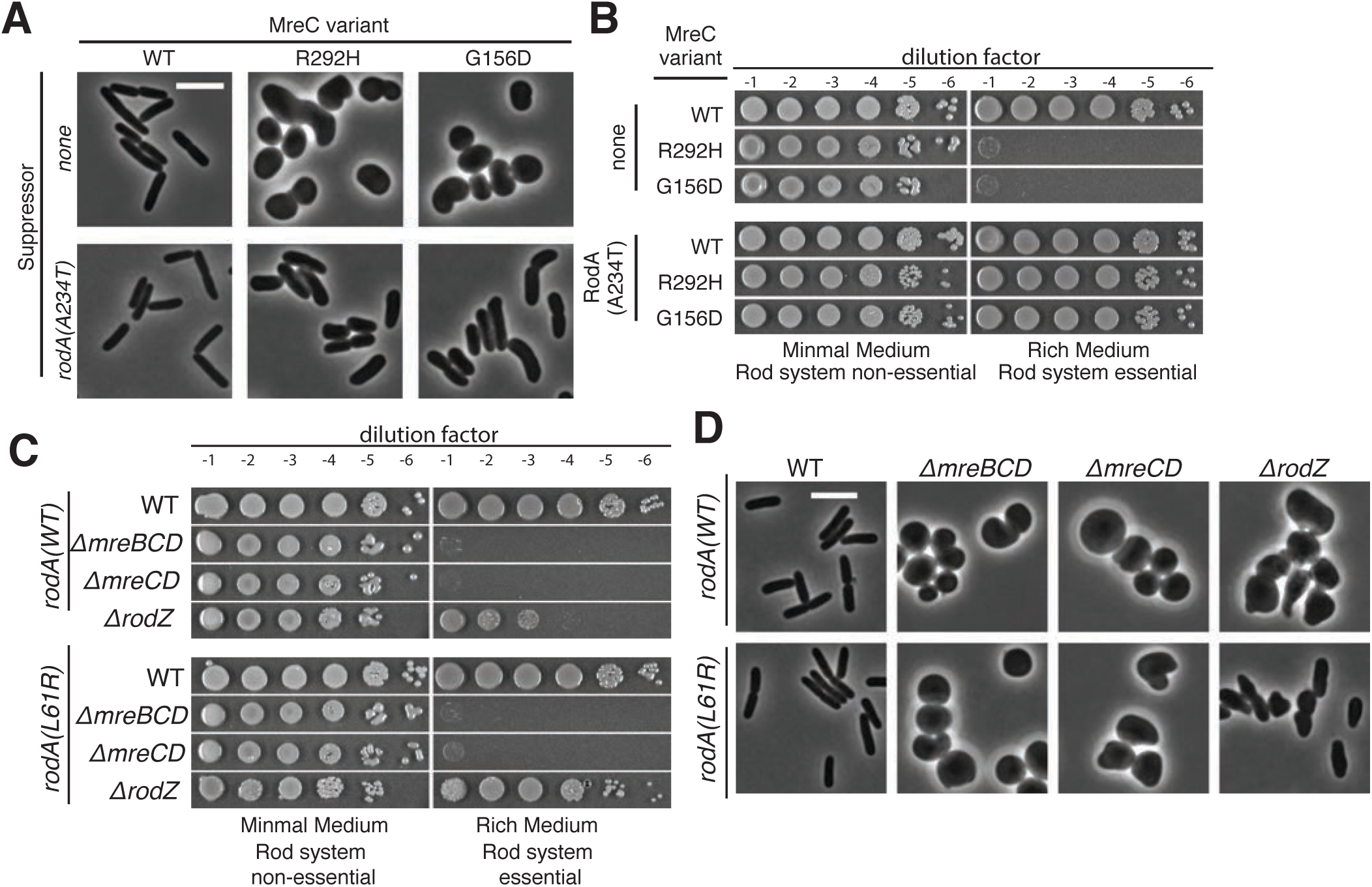
RodA(A234T) suppresses *mreC* point mutants and *ΔrodZ* but not Δ*mreCD*. **A.** Strains containing the indicated point mutations at the native genomic locus [PR158, PR159, PR160, PR161, PR162, PR163] were grown and imaged as in Figure 2A. **B.** Overnight cultures of the above strains were serially diluted and spotted on either M9 CAA glu agar (Rod non-essential) or LB agar (Rod essential). Plates were incubated at 30℃ for either 40 h (M9) or 16 h (LB) before imaging. **C.** Overnight cultures of the indicated strains [PR150, PR152, PR153, PR154, PR151, PR155, PR156, PR157] were were serially diluted and spotted as in Figure 3B. **D.** The indicated strains were grown, fixed, and imaged as described in Figure 2A.

The ability of RodA and PBP2 variants to stimulate PG synthesis by the Rod system suggested that activation in both cases may ultimately result from the enhancement of PG polymerization by RodA. To test this possibility more directly, we used a modified radiolabeling assay in which the beta-lactam mecillinam was included. Mecillinam specifically blocks the TP activity of PBP2 but allows continued glycan polymerization by RodA (Cho et al., 2016). We previously showed that the uncrosslinked glycans produced in mecillinam-treated cells are rapidly degraded by the lytic transglycosylase Slt to form soluble turnover products (anhydromuropeptides) (Cho et al., 2014). Thus, in radiolabeled cells simultaneously inhibited for cell division and treated with mecillinam, the level of labeled turnover products produced provides a measure of RodA polymerization activity (Figure 3D). Using this assay, we found that both RodA(A234T) and PBP2(L61R) resulted in elevated PG turnover in mecillinam treated cells (Figure 3E-F). Similar assays were performed to monitor the effects of Rod system variants on aPBP activity using the beta-lactam cefsulodin. This antibiotic specifically inhibits the transpeptidase activity of aPBPs such that PG turnover in cefsulodin-treated cells provides a measure of aPBP PG polymerase activity (Cho et al., 2014; 2016). Cefsulodin-induced PG turnover was found to be reduced in both RodA(A234T) and PBP2(L61R) containing cells (Figure 3, supplement 2), indicating a reduction of aPBP polymerase activity. This reduction in activity most likely reflects an increased competition for precursors between aPBPs and the activated Rod system. Based on the radiolabeling results we conclude that the RodA(A234T) and PBP2(L61R) variants enhance Rod system function by promoting PG polymerization by RodA.

### PBP2(L61R) activates PG polymerization by RodA in purified RodA-PBP2 complexes

The in vivo labeling results suggest the attractive possibility that changes in PBP2 structure, either through its interaction with MreC or the L61R substitution, can be communicated to RodA to activate PG polymerization. We therefore wanted to test this potential RodA activation mechanism in vitro using purified RodA-PBP2 complexes. To simplify purification of the complexes, we generated a RodA-PBP2 fusion protein with the two components connected by a linker (GGGSx3). A similar SEDS-bPBP fusion had been shown to be functional for *Bacillus subtilis* sporulation (Fay et al., 2010). Our construct was also active in vivo as it largely restored rod shape to Δ*pbpA-rodA* cells (Figure 5, supplement 1). We therefore proceeded to purify a FLAG-tagged version of the wild-type fusion and fusions harboring either PBP2(L61R) or RodA(A234T). The fusions were produced in an *E. coli* expression strain lacking three of its four aPBP-type PG polymerases (PBP1b, PBP1c, and MtgA) to limit the potential for contaminating polymerase activity in the purified preparations. The resulting preparations were >90% pure with some observable lower molecular weight material. We suspect that most of this material is derived from cleavage of the fusion within the linker as the bands migrate at ∼70 kDa and ∼40 kDa corresponding to the molecular weights of PBP2 and RodA, respectively (Figure 5A).

**Figure 5.**
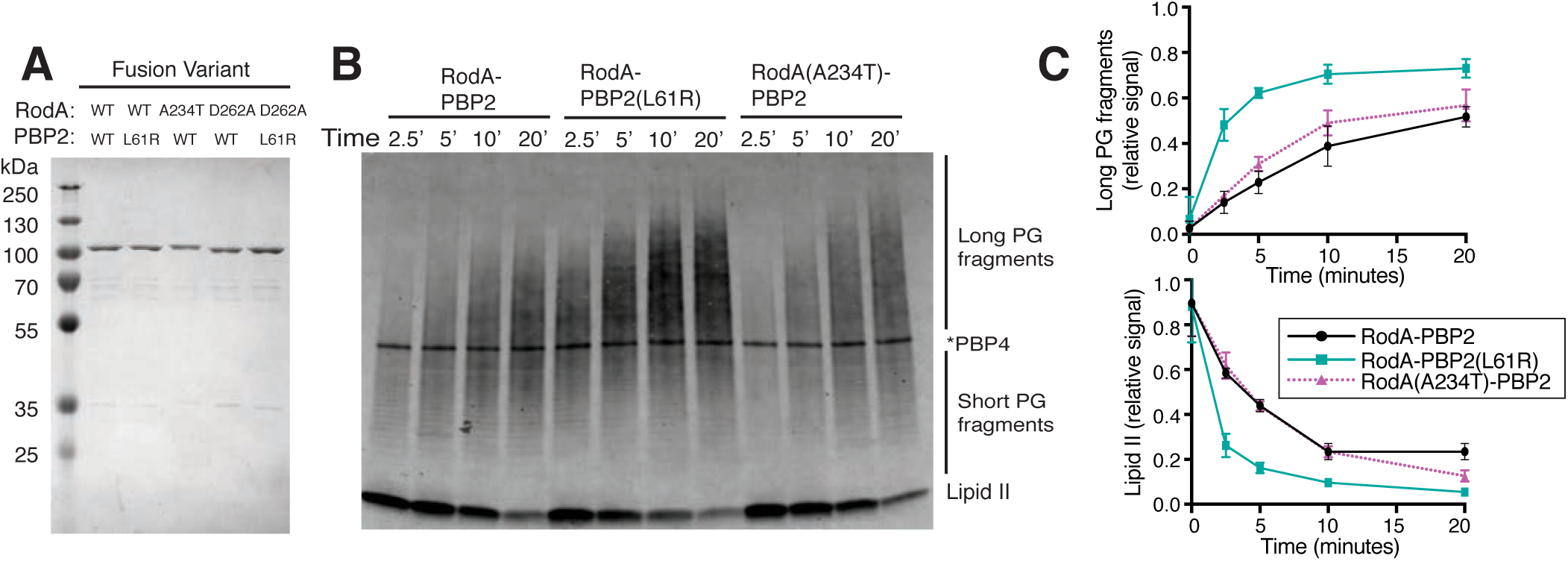
PBP2(L61R) stimulates glycosyltransferase activity of RodA. **A.** Purified Flag-RodA-PBP2 and mutant derivatives were run on a Coomassie-stained SDS-PAGE gel. The molecular weight of the fusion proteins is approximately 114 kDa. **B.** Blot detecting the peptidoglycan product produced by the RodA-PBP2 fusion constructs incubated with extracted *E. coli* Lipid II for the indicated length of time. The product was detected by BDL labeling with *S. aureus* PBP4, which appears as a labeled band in the middle of the blot. **D.** The accumulation of long PG fragments and depletion of lipid II during three independent replicates of glycosyltransferase time-courses were quantified using densitometry. Error bars represent standard deviation.

We first compared the polymerase activity of RodA-PBP2(WT) with RodA-PBP2(L61R) and RodA(A234T)-PBP2. Purified lipid II substrate from *E. coli* was added to the fusions and the reactions terminated at various time points following initiation. The resulting products were then subjected to enzymatic labeling with biotin-D-lysine, separated on an SDS-PAGE gel, transferred to a PVDF membrane, and detected with streptavidin conjugated to an infrared dye (Qiao et al., 2017). Mecillinam was included in the reactions to prevent glycan crosslinking by PBP2 so that polymer length could be determined without complications from crosslinking by PBP2. All fusions promoted the production of glycan polymers that increased in abundance and apparent length over time (Figure 5B, C). However, the RodA-PBP2(L61R) generated product more rapidly than RodA-PBP2(WT) and produced products that were longer (Figure 5B, C). The length and amount of PG produced by RodA(A234T)-PBP2 was not statistically different than the wild-type fusion (Figure 5B, C). Notably, the polymerase activity of all fusions was insensitive to moenomycin, an inhibitor that blocks aPBP-type PGT activity (Figure 5, supplement 2). Also, the polymerase activity of fusions with PBP2(WT) and PBP2(L61R) was completely blocked by a D262A substitution in RodA (Figure 5, supplement 2). An equivalent change was previously shown to inactivate the polymerase activity of *B. subtilis* RodA (Meeske et al., 2016). Therefore, the polymerase activity observed for the fusions is unlikely to be due to contaminating PBP1a, the only aPBP-type polymerase produced in the expression strain. We conclude that SEDS-bPBP complexes indeed form a functional PG synthase as proposed previously (Cho et al., 2016; Meeske et al., 2016), and that changes in the bPBP can be communicated to the SEDS protein to stimulate its PG polymerase activity.

### PBP2(L61R) increases the number of functional Rod complexes per cell

Fluorescent protein fusions to MreB and other Rod system components in *E. coli* and *B. subtilis* form multiple dynamic foci dispersed throughout the cell cylinder. These foci have been observed to rotate in a processive manner around the long axis of the cell (Cho et al., 2016; Domínguez-Escobar et al., 2011; Garner et al., 2011; van Teeffelen et al., 2011), and this motion is blocked by inhibitors of PG synthesis. Thus, the dynamic behavior of MreB and other Rod components is thought to be driven by the deposition of new PG material into the matrix with the speed of rotational movement reflecting the synthetic activity of the Rod complex.

To further understand the mechanism of Rod system activation by the PBP2(L61R) variant, we monitored its effect on the localization dynamics of MreB and PBP2 using total internal reflection fluorescence (TIRF) microscopy. An MreB sandwich fusion with mNeonGreen (^SW^MreB-mNeon) and an N-terminal superfolder-GFP fusion to PBP2 (sfGFP-PBP2) were used for the imaging. Both fusions were previously shown to be functional (Cho et al., 2016). ^SW^MreB-mNeon foci displayed processive rotational movement in cells producing PBP2(WT) or PBP2(L61R) (**Movie S1-2**). The speed of rotational movement was unchanged by the PBP2(L61R) variant (Figure 6A,C). Similarly, sfGFP-PBP2(WT) and sfGFP-PBP2(L61R) formed foci that moved around the cell long axis with a almost identical velocities (**Movie S3-4**, Figure 6B-C). Although the speed of particle motion was unchanged by the PBP2(L61R) variant in each case, the number of moving particles per cell appeared to increase in cells producing the altered PBP2. We therefore quantified the number of particle tracks per cell for each imaging experiment. Indeed, more directionally moving ^SW^MreB-mNeon foci were observed per cell in the PBP2(L61R) producing cells versus those with PBP2(WT) (Figure 6D). Likewise, cells expressing sfGFP-PBP2(L61R) possessed a greater number of directionally moving foci than those producing sfGFP-PBP2(WT) (Figure 6E). These results suggest that PBP2(L61R) not only stimulates RodA polymerase activity, but also promotes the assembly of more active Rod complexes per cell.

**Figure 6.**
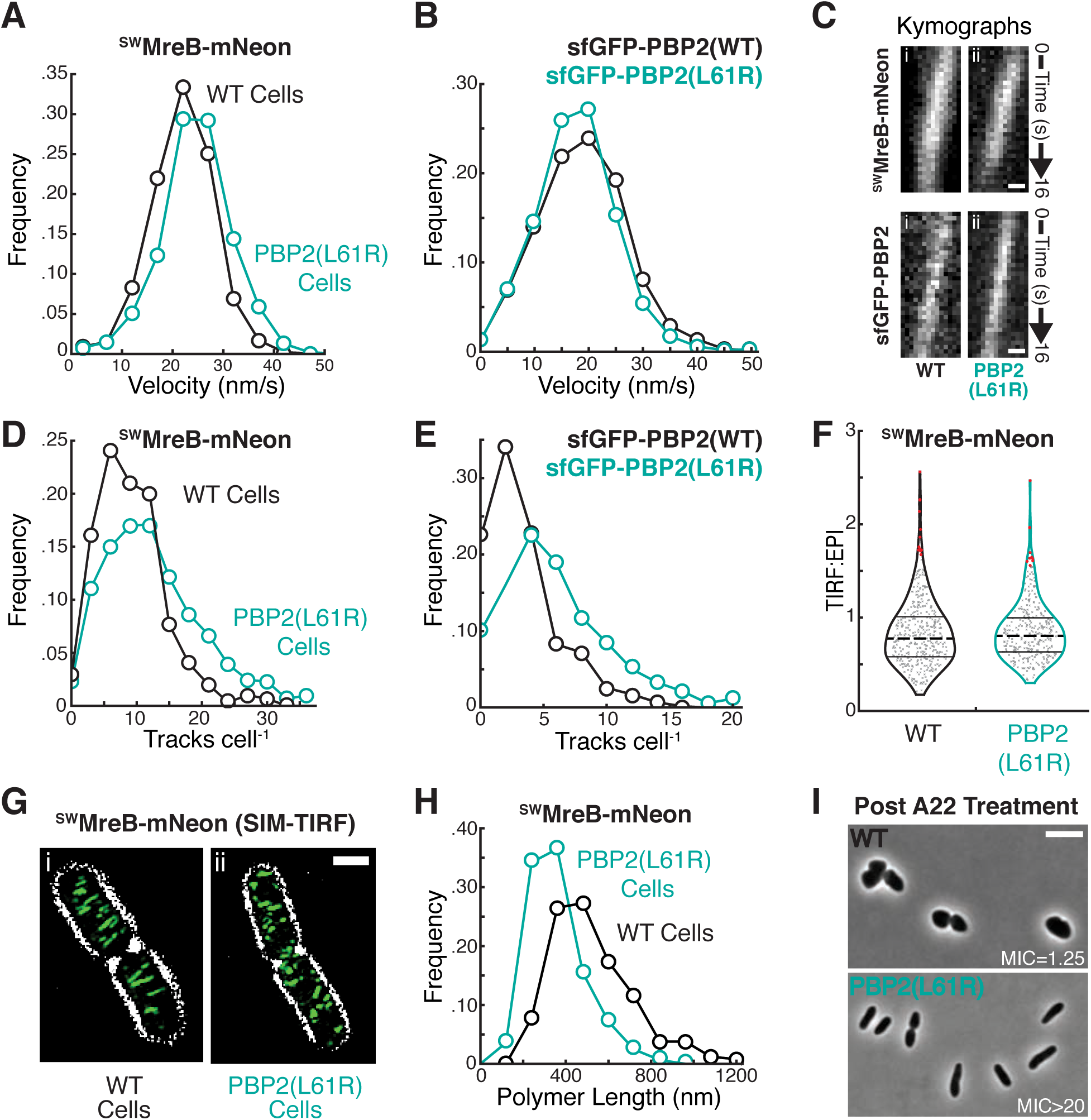
Rod system dynamics in PBP2(L61R) cells. **A.** Histograms of velocity measurements determined for individual traces of ^SW^MreB-mNeon (attλHC897) in wild-type (22±6 nm/s, n=1467; black) and PBP2(L61R) cells [PR78] (25±7 nm/s, n=949; turquoise). **B.** Histograms of velocity measurements determined for individual tracks of sfGFP-PBP2 (attλHC943) (18.9±8.1 nm/s, n=2692; black) and sfGFP-PBP2(L61R) (attλPR128) (17.8±7.5 nm/s, n=3440; turquoise) in Δ*pbpA* cells. Bin size, 5 m/s. **C.** Kymographs of individual ^SW^MreB-mNeon (attλHC897) tracks in wild-type (i) and PBP2(L61R) (ii) cells are displayed atop kymographs of individual sfGFP-PBP2 (attλHC943, i) or sfGFP-PBP2(L61R) (attλPR128, ii) tracks in Δ*pbpA* cells. Scale bar, 250 nm. **D.** Histograms for the number of ^SW^MreB-mNeon tracks measured per cell in wild-type (9.0±5.3, n=609; black) and PBP2(L61R) cells (12.4±7.5, n=300; turquoise) 37℃. **E.** Histograms for the number of sfGFP-PBP2 (3.8±3.0, n=581; black) and sfGFP-PBP2 (6.4±4.2, n=517; turquoise) tracks measured per cell in Δ*pbpA* cells at 30℃. As expected (Billaudeau et al., 2017) there are less directionally moving rod complexes at lower temperatures. Bin size, 3 tracks cell^-1^. **F.** Violin plots illustrating the distribution of normalized fluorescence measurements for ^SW^MreB-mNeon (attλHC897) expressed in wild-type (0.83±0.36, n=397) and PBP2(L61R) cells (0.85±0.31, n=321). The fluorescence intensity acquired under TIRF illumination for individual cells was integrated and divided by similar measurements taken under EPI illumination, providing an approximation for the relative abundance of surface-associated ^SW^MreB-mNeon. The distribution of values along the x-axis capture the frequency of measurements along the y-axis. Lines designate quartiles with the dotted line indicating the mean value. Outliers are highlighted in red. **G.** Representative SIM-TIRF micrographs of ^SW^MreB-mNeon integrated at the native locus in wild-type [JAB593] and PBP2(L61R) cells [JAB576]. The signal for ^SW^MreB-mNeon is pseudocolored green and overlaid a contrast-adjusted phase-contrast image. Scale bar, 1μm. **H.** Distributions of ^SW^MreB-mNeon polymer lengths in wild-type [JAB593] (520±190 nm, n=502; black) and PBP2(L62R) cells [JAB576] (360±130 nm, n=614; turquoise). Bin size, 120 nm. **I.** Representative micrographs of wild-type [MG1655] or PBP2(L61R) [PR78] cells after a 4hr treatment with 2 μg/mL A22, an MreB-inhibitor. The minimum inhibitory concentration (MIC) of A22 for each cell type is displayed in μg/mL.

One possible way in which the PBP2(L61R) variant could increase the number of active Rod complexes per cell is via enhancing the recruitment of MreB filaments to the membrane. To investigate this possibility, we measured the total ^SW^MreB-mNeon fluorescence per cell by epifluorescence (EPI) illumination and the fluorescence at the cell surface using TIRF illumination. We then calculated the TIRF/EPI ratio for each cell as a measure of MreB membrane recruitment. To ensure equivalent illumination of cells producing PBP2(WT) or PBP2(L61R), we introduced a cytoplasmic mCherry marker into one of the strains, mixed them, and performed the TIRF and EPI measurements on both strains simultaneously. Strain identity was then determined by the presence or absence of the mCherry marker (Figure 6, supplement 1). Two sets of measurements were made, one with the marked strain being PBP2(WT) and the other with the PBP2(L61R) strain being marked. The analysis revealed no significant change in the TIRF/EPI ratio of ^SW^MreB-mNeon fluorescence between cells with either PBP2(WT) or PBP2(L61R) (Figure 6F), indicating that the total amount of MreB recruited to the membrane is not altered by PBP2(L61R).

The observation that the PBP2(L61R) variant increases the number of directionally moving ^SW^MreB-mNeon foci per cell without increasing the total amount of MreB at the membrane suggested that the altered synthase may be modulating MreB filament formation. To investigate this possibility, we imaged ^SW^MreB-mNeon using structured-illumination microscopy combined with TIRF illumination (SIM-TIRF). With this super-resolution method, clear filaments of ^SW^MreB-mNeon were visible that displayed a dynamic circumferential motion like the foci observed at lower resolution (Figure 6G, **Movie S5**). Analysis of still images of cells with PBP2(WT) or PBP2(L61R) allowed us to measure the differences in length of the fluorescent MreB filaments. Strikingly, the filaments observed in PBP2(L61R) cells were on average significantly shorter than those found in cells producing PBP2(WT) (Figure 6H). This observation suggests that changes in PBP2 affect MreB polymer formation and/or dynamics. Accordingly, similar to previously isolated PBP2 and RodA variants, cells producing PBP2(L61R) are resistant to the MreB antagonist A22 (Figure 6I), indicating that MreB polymers are more robust in these cells. Overall, the cytological results are consistent with a model in which the activation status of the core PG synthase of the Rod system is communicated to MreB to control filament formation so that the new synthesis promoted by the activated enzymes is properly oriented.

## DISCUSSION

Cell shape determination in bacteria requires control of when and where new PG is made and incorporated into the existing matrix. It has been clear for some time that this spatiotemporal regulation is mediated by multiprotein complexes linked to cytoskeletal filaments (Typas et al., 2012). However, an understanding of how the PG synthase enzymes within these machines are regulated has been lacking. It has also remained unclear how the polymerization state of the cytoskeletal filaments might affect the activation status of the synthases or vice versa. Our investigation of Rod system function suggests that its activity is governed by an activation pathway involving components of the machinery with heretofore unknown function (Figure 7). The results also provide insight into how the synthetic activities of the PG polymerase and crosslinking enzyme within the complex are coordinated. Moreover, our results support a model in which activated PG synthesis enzymes exert control over MreB polymer formation, suggesting that MreB polymerization does not serve as the primary regulatory step in Rod system activation. Finally, based on the similar nature of mutants activated for Rod system function to those bypassing normal regulation of the division machinery, we propose that all morphogenic machines are likely to be governed by an activation pathway controlling SEDS-bPBP synthases analogous to the one described here for Rod system regulation.

**Figure 7:**
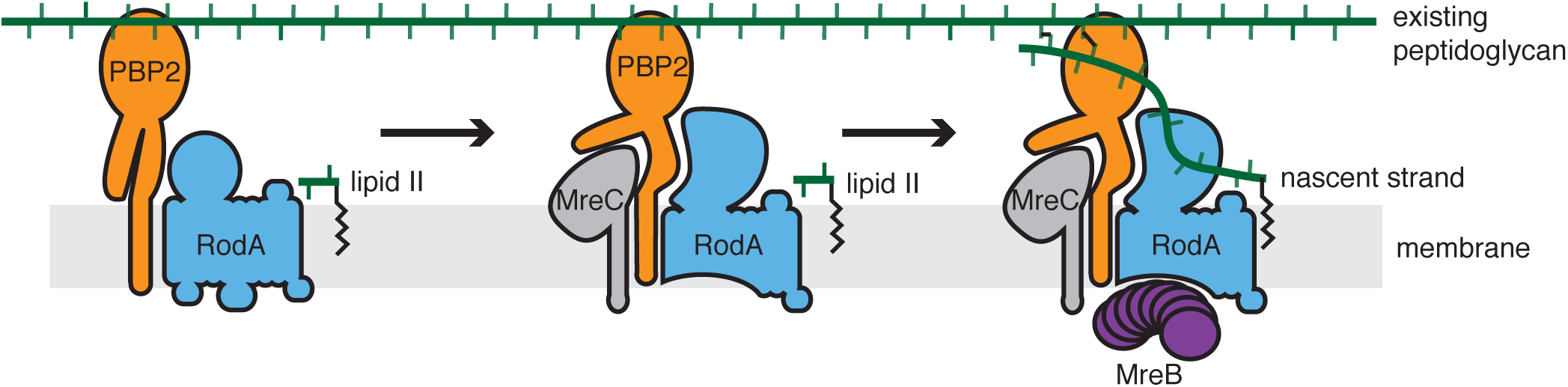
Proposed activation pathway governing peptidoglycan synthesis by the Rod System. In the absence of other factors, PBP2 and RodA are enzymatically inactive (left). In response to signals yet to be determined, MreC associates with PBP2, induced a conformational change, which in turn activates RodA (center). We propose that this activated complex has a higher affinity for MreB to promote MreB filament formation (right) to orient the PG produced by the activated synthase. For simplicity, MreD and RodZ are not shown but are also required for proper Rod system function, possibly by modulating the MreC-PBP2 interaction.

### A potential activation pathway controlling Rod system function

To gain insight into the regulation of the Rod system, we selected for suppressors of *mreC* point mutants. Although the precise nature of the functional defect(s) caused by these mutations remains to be determined, they allowed us to identify two PBP2 variants that activate the Rod system. This activation both bypasses the need for some Rod system proteins, and hyperactivates the Rod system in otherwise wild-type cells. Characterization of the suppressor mutants combined with a recently solved structure of an MreC-PBP2 complex from *H. pylori* (Contreras-Martel et al., 2017) supports a regulatory role for MreC in Rod system activation.

In the structure of the MreC-PBP2 complex, MreC was found to induce a significant conformational change in the membrane-proximal pedestal domain of PBP2, causing two of its subdomains to hinge open (Figure 2C) (Contreras-Martel et al., 2017). The amino-acid changes in PBP2 that suppress MreC defects mapped to the same region of the protein, suggesting that they may promote a conformation of PBP2 that mimics that induced by MreC. Biochemical and physiological results indicate that one of these altered PBP2 proteins, PBP2(L61R), not only suppresses MreC defects, it also stimulates Rod system activity in vivo and PG synthesis by RodA-PBP2 fusions in vitro. We infer from the combined set of results that the interaction between PBP2 and MreC is probably not just a scaffolding interaction as proposed previously (Contreras-Martel et al., 2017), but also likely serves a regulatory role in Rod system function by shifting the RodA-PBP2 PG synthase into an activated conformation (Figure 7). Although a direct role for MreC in promoting RodA-PBP2 synthase activity remains to be tested, such an activation mechanism would ensure that the PG synthase is only highly active in the context of the assembled Rod complex thereby providing spatiotemporal control over its function.

In addition to suppressing the Rod system defect caused by missense alleles of *mreC*, the PBP2(L61R) variant also promoted viability and partially restored rod-shape to mutants deleted for *mreC*, *mreCD*, and *rodZ* as well as a triple *mreCD rodZ* deletion. However, the same PBP2 variant failed to suppress an *mreBCD* deletion, indicating that MreB is needed for Rod system function even when the core enzymes are abnormally activated. This MreB-requirement most likely reflects the important role of MreB filaments in promoting rod-shape by orienting the motion of the synthetic enzymes (Hussain et al., 2018). In this regard, the ability of PBP2(L61R) to promote partial Rod system function in the triple *mreCD rodZ* deletion is remarkable because it implies that MreB can interface directly with the RodA-PBP2 synthase. Thus, a cytoskeletal filament connected to a PG synthase complex appears to be the minimal functional unit of the Rod system. The other components of the system are likely to be important for stabilizing the connection between RodA-PBP2 and MreB. However, because MreC, MreD, and RodZ are conserved along with RodA and PBP2 in ovoid and spherical bacteria lacking MreB, it seems unlikely that their sole function is to provide bridging interactions between the enzymes and MreB filaments. Instead, this conservation in combination with the suppression results with PBP2(L61R) suggests that like MreC, MreD and RodZ are probably also involved in promoting the activation of PG synthesis by RodA-PBP2, either directly or through an effect on the MreC-PBP2 interaction.

### Control of MreB polymerization by the activation status of the RodA-PBP2 synthase

PBP2(L61R) cells were found to assemble more circumferentially moving MreB and PBP2 foci than PBP2(WT) cells. Additionally, super-resolution microscopy revealed that the MreB filaments formed at the membrane were shorter in the cells with the activated PBP2 variant. An increase in polymer number with a corresponding decrease in length is expected if polymer formation is stimulated without a change in the monomer supply. PBP2(L61R) was not found to alter the cellular MreB concentration or the total amount of MreB recruited to the membrane (**Figure 2 supplement 1**, Figure 6 supplement 1). Thus, the cytological results support a role for RodA-PBP2 activation in enhancing MreB polymerization, potentially by nucleating the formation of new polymers, either directly or through effects of the activated synthase on other Rod system components like RodZ (Morgenstein, Bratton, Shaevitz, & Gitai, 2017). Another connection between RodA-PBP2 activation and MreB polymerization comes from the observation that PBP2(L61R), and previously isolated PBP2 and RodA variants that are presumably also activated, confer resistance to the MreB antagonist A22 (Figure 6I) (Shiomi et al., 2013), indicating that they somehow making polymer formation more robust to disruption by the drug. Finally, MreB filament formation at the membrane has previously been shown to be dependent on the availability of the RodA-PBP2 substrate lipid II in *B. subtilis* (Schirner et al., 2015). Taken together, these observations support a model in which factors upstream of MreB polymerization are important control points in Rod system assembly and activation. Given the regulatory roles for MreC, MreD, and RodZ implied by the genetic results, an attractive possibility is that the membrane and periplasmic domains of these proteins function as sensors that promote PG synthesis by the Rod system in response to chemical and/or physical signals from the cell envelope such as PG crosslinking status, membrane curvature, or physical strain (Ursell et al., 2014; Wong et al., 2017). In this scenario, MreB filaments would be polymerized at or recruited to sites where synthesis is activated by the membrane-embedded components. Once recruited, MreB could then act as a rudder to steer cell wall insertion along the circumferential axis (Hussain et al., 2018). It is also possible that the activation process is initiated by MreB polymerization induced by a different set of stimuli. Importantly, the two possibilities are not mutually exclusive, and it may well be that multiple inputs into the formation of active Rod complexes contribute to the robustness of the system in promoting rod shape. A major challenge moving forward will be to determine the molecular nature of the signals to which the Rod system is responding to trigger its synthetic activity.

### Coupling of PG polymerization and crosslinking within the Rod system

Complexes between SEDS and bPBPs have been well described for the divisome (FtsW-PBP3) and sporulation (SpoVE-SpoVD) (Fay et al., 2010; Fraipont et al., 2011). Therefore, following the discovery of PG polymerase activity for RodA, it was proposed that RodA-PBP2 and other SEDS-bPBP complexes form a functional PG synthase with both polymerase and crosslinking activity (Cho et al., 2016; Meeske et al., 2016). This possibility is supported by recent evolutionary coupling analyses and mutational studies indicating that a RodA-PBP2 complex formed through interactions between RodA and the pedestal domain of PBP2 is likely to be critical for Rod system function (Sjodt et al., 2018). Here, we found that changes in the PBP2 pedestal domain can activate PG synthesis by RodA in vivo and stimulate the activity of RodA-PBP2 fusions in vitro. Together, these observations suggest that the RodA-PBP2 complex not only physically connects the two enzymes, but also serves as a regulatory conduit used to coordinate their activities. In this case, the genetic, biochemical, and structural data support a model in which conformational changes in the pedestal domain of PBP2 induced by MreC, likely in conjunction with other components of the system, are communicated to RodA to stimulate PG synthesis. This level of communication between the PGT and TP enzymes is attractive because it would provide a means to prevent RodA from robustly producing glycan strands without the ability to crosslink them. Otherwise, as revealed by experiments with the beta-lactam mecillinam, the production of uncrosslinked glycans by RodA when PBP2 is inactive results in a toxic futile cycle of glycan synthesis and degradation (Cho et al., 2014).

### A possible conserved regulatory mechanism governing PG synthesis by SEDS-bPBP synthases

Based on analogy with RodA-PBP2, FtsW-PBP3 has been proposed to be the core PG synthase of the divisome (Cho et al., 2016; Meeske et al., 2016). Recent biochemical studies from our laboratories indicate that FtsW indeed possesses PG polymerase activity and that this activity requires the formation of a complex with its cognate bPBP (Taguchi et al., 2018). This finding is consistent with a required coupling between PG polymerase and crosslinking functions to prevent the formation of toxic uncrosslinked glycans. Genetic evidence in the literature also suggests that the FtsW-PBP3 complex is regulated by a mechanism analogous to that of RodA-PBP2. Several gain-of-function alleles in the genes encoding FtsW and PBP3 were previously isolated as suppressors of division inhibitor overproduction in *Caulobacter cresentus* and *E. coli* (Du et al., 2016; Modell et al., 2011; 2014).

Notably, FtsW(A246T) was one of the suppressors of division inhibition identified in *C. cresentus* (Modell et al., 2014). This residue change corresponds to A234T in *E. coli* RodA, the exact change that we and others have found to activate PG biogenesis by the Rod system and suppresses defects in MreC and RodZ (Shiomi et al., 2013). Moreover, the amino acid substitutions in PBP3 that suppress division inhibition in *C. cresentus* map to the N-terminal domain not far from where we have found alterations in PBP2 that hyperactivate the Rod system (Modell et al., 2011). Thus, the genetic evidence points towards PG biogenesis by the divisome being activated by the FtsW and PBP3 variants such that normal regulatory controls governing the activity of the complex can be bypassed. The similarity of these changes to those in RodA and PBP2 that activate the Rod system suggest that SEDS-bPBP complexes within morphogenic machines are likely to be regulated by similar and broadly conserved mechanisms. This activation step therefore represents an attractive target for small molecule inhibitors for use in antibiotic development.

## MATERIALS AND METHODS

### Media, bacterial strains, and plasmids

All *E. coli* strains used in the reported experiments are derivatives of MG1655 (Guyer, Reed, Steitz, & Low, 1981). Strains were grown in LB (1% tryptone, 0.5% yeast extract, 0.5% NaCl) or minimal M9 medium (J. H. Miller, 1972) supplemented with 0.2% casamino acids and 0.2% glucose (abbreviated M9 CAA glu). Unless otherwise indicated, antibiotics were used at 25 (chloramphenicol; Cm), 50 (kanamycin; Kan), 50 (ampicillin; Amp), 50 (spectinomycin; Spec), or 5 (tetracycline; Tet) μg/mL. Growth conditions for microscopy experiments are described in the figure legends.

### Molecular biology

PCR was performed using Q5 polymerase (NEB) according to the manufacturer’s instructions. Plasmid DNA and PCR fragments were purified using the Zyppy plasmid miniprep kit (Zymo Research) or the Qiaquick PCR purification kit (Qiagen), respectively. Sequencing reactions were carried out with an ABI3730xl DNA analyzer at the DNA Resource Core of Dana-Farber/Harvard Cancer Center (funded in part by NCI Cancer Center support grant 2P30CA006516-48).

### Selection for suppressors of *mreC* point mutants

Overnight cultures of PR5 [*mreC(R292H)*] or PR30 [*mreC(G156D)*] were grown at 30℃ in M9 medium supplemented with casamino acids and glucose. Serial dilutions of these cultures were plated on both permissive conditions (M9 CAA glu agar at 30℃) and conditions that are non-permissive for the growth and survival of spherical cells (LB supplemented with 1% sodium dodecyl sulfate (SDS) at 30℃ or 37℃) (Bendezú et al., 2009). After 24 hours of incubation colonies that appeared on the LB + SDS plates were replica streaked on LB agar and LB agar supplemented with 10 μg/mL A22. We reasoned that suppressor mutants that have restored rod system function would be sensitive to A22 (A22^S^), whereas mutants that had found an alternative means to survive on LB, such as overexpression of *ftsZ,* would be resistant to A22 (A22^R^). All SDS^R^, A22^S^ isolates were visually screened for restoration of rod cell shape using a Nikon Eclipse 50i microscope equipped with a 100x Ph3 DL 1.25 NA lens (rig #3, see below). Overnight liquid cultures of SDS^R^, A22^S^, rod shaped isolates were grown in LB at 30℃, and genomic DNA was prepared using a Wizard^®^ Genomic DNA Purification Kit (Promega) and Genomic DNA Clean & Concentrator™-10 (Zymo Research).

Two different methods were used for whole genome sequencing of suppressor strains. Some suppressors were prepared for sequencing using a modified Nextera library preparation strategy, as described by Baym et al. (Baym et al., 2015). Other suppressors were prepared for sequencing using the NEBNext^®^ Ultra™ DNA Library Prep Kit for Illumina^®^ according to manufacturer’s instructions. DNA concentrations were determined using the Qubit^®^ dsDNA HS Assay Kit and sizes were determined using a High Sensitivity D1000 screen tape run on an Agilent 4200 TapeStation system. Sequencing was performed using a MiSeq Reagent Kit v3, with the Miseq System (Illumina). Reads were mapped using the CLC Genomics Workbench software (Qiagen).

### Immunoblotting

Proteins were run on a 10% SDS-PAGE gel and transferred to an activated PVDF membrane. The membrane was briefly rinsed, then blocked with 2% milk (w/v) in Tris-buffered saline, 0.1% Tween-20 (TBS-T) for 1 hour at room temperature. The membrane was then transferred to primary antibody solution, containing 0.2% milk (w/v), rabbit anti-MreB (Bendezú et al., 2009) or rabbit anti-MreC (1:10,000 dilution) and mouse anti-RpoA (BioLegend clone 4RA2, 1:10000 dilution) in TBS-T, and incubated for 16 hours at 4℃. The membrane was rinsed quickly, then washed three times for ten minutes in TBS-T. The membrane was transferred to a solution of secondary antibodies (anti-rabbit 800CW and anti-mouse 680RD; Li-COR) in 0.1% milk for 1 hour at room temperature. After four ten-minute washes in TBS-T, the membrane was imaged using a Li-COR ODESSEY Clx scanner.

**Bocillin binding assays** were performed as described previously (Cho et al., 2016) **^3^H-mDAP physiological radiolabeling-**Peptidoglycan precursor levels, synthesis, and turnover were determined as described previously (Cho et al., 2014; 2016). The results were analyzed using a two-way ANOVA, followed by Tukey’s multiple comparisons test.

### Protein expression

The strain used for expression of the RodA-PBP2 fusions was an *E. coli* C43 derivative of BL21(DE3) with deletions in *ponB*, *pbpC*, *mtgA* (strain CAM333) that contains a plasmid expressing Ulp1 (403-621) protease under an arabinose-inducible promoter (pAM174) (Meeske et al., 2016). The RodA-GGGSx3-PBP2 fusion constructs were overexpressed with a His6-SUMO-Flag tag fused to the N-terminus, as described previously (Meeske et al., 2016). The plasmids bearing the protein fusion (pSS50) and mutant derivatives (pSS51, pSS52, pSS60, pSS62) were transformed into CAM333 under antibiotic selection. Transformants for each construct were used to inoculate 5 mL of LB media supplemented with ampicillin (50 μg/ml) and chloramphenicol (25 μg/mL) and were grown overnight at 37℃. Cultures were then diluted into 1L of Terrific Broth medium, supplemented with 0.1% glucose and 2 mM MgCl_2_, and grown at 37℃ to OD_600_ of 0.8. IPTG was then added to 1 mM to induce expression of the fusion, and arabinose was added to 0.1% to induce expression of Ulp1. After induction overnight at 20℃, the cells were harvested by centrifugation. The cell pellets were resuspended in lysis buffer (50 mM HEPES pH 7.5, 150 mM NaCl, 20 mM MgCl_2_, 0.5 M DTT) and lysed by passage through a french press twice at 25,000 psi. Membranes were collected by ultracentrifugation at 100,000g for 1 hour at 4℃. The membrane pellets were mechanically homogenized by a teflon dounce and solubilized in buffer containing 20 mM HEPES pH 7.0, 0.5 M NaCl, 20% glycerol, and 1% n-dodecyl-B-D-maltoside (DDM) for 2 hours at 4℃. Insoluble material was pelleted by ultracentrifugation at 100,000g for 1 hour at 4℃. The soluble fraction was removed and supplemented with 2 mM CaCl_2_ and applied to homemade M1 anti-Flag antibody resin. The resin was washed with 25 mL of wash buffer (20 mM HEPES pH 7.0, 0.5 M NaCl, 20% glycerol, 2 mM CaCl_2_, 0.1% DDM). The Flag-tagged constructs were eluted from the resin in 1 mL fractions with buffer containing 20 mM HEPES pH 7.0, 0.5 M NaCl, 20% glycerol, 0.1% DDM, 5 mM EDTA pH 8.0, and 0.2 mg/mL 3X FLAG peptide (Sigma). The purity of the sample was checked by SDS-PAGE. The final yield for each of the different fusion constructs was approximately 1 mg per 1 L of culture.

A His-SUMO tagged version of the soluble domain of MreC (amino acids 45-367) was purified and used for antibody production. Lemo21(λDE3)/pPR57 cells were grown in LB supplemented with 5 mg/mL ampicillin and 25 mg/mL chloramphenicol and grown at 37℃ until the OD_600_ reached 0.4. Cells were then induced with 1 mM IPTG and grown for an additional 2 hours. Cells were pelleted and resuspended in buffer A (20 mM Tris-HCl (pH=8.0), 300 mM NaCl, 0.5 mM DTT, 20% glycerol) containing 30 mM imidazole. cells were disrupted by passing them through a french pressure cell twice at 15,000 psi. Cell debris and membranes were pelleted by centrifugation at 100,000 x g for 30 minutes at 4℃. The resulting extract was mixed with pre-equilibrated QIAGEN Ni-NTA agarose beads, then transferred to a column. The column was washed sequentially with buffer A containing 30 mM, 50 mM, and 100 mM imidazole, then eluted in buffer A containing 300 mM imidazole. The eluate was digested with His-Ulp1 to cleave the His-SUMO tag, dialyzed in buffer A, then run through the Ni-NTA column to obtain pure, untagged MreC. Purified protein was sent to Covance Inc. for the production of rabbit polyclonal antibodies.

### Peptidoglycan glycosyltransferase activity assay

Purified proteins were concentrated to 10 μM using a 100 kDa MWCO Amicon Ultra Centrifugal Filter (Millipore). Extraction of *E. coli* Lipid II was performed as described previously (Qiao et al., 2017). Peptidoglycan glycosyltransferase activity was assayed as previously described (Srisuknimit et al., 2017). Briefly, Lipid II dissolved in DMSO (2 μM) was incubated with each purified protein (1 μM) with 1X reaction buffer in a total volume of 10 μL for 20 minutes at room temperature, unless otherwise indicated. The reaction buffer contains 50 mM HEPES pH 7.0, 20 mM MgCl_2_, 20 mM CaCl_2_, 200 μM mecillinam, and 20% DMSO. Moenomycin dissolved in DMSO was used at a final concentration of 3 μM. Reactions were quenched by incubation at 95℃ for 2 minutes. Biotinylation of the peptidoglycan product was subsequently performed by addition of 2 μL of 20 mM Biotin D-Lysine (BDL) and 1 μL of 50 μM *S. aureus* PBP4 (Kahne lab) and incubation at room temperature for 1 hour. The reaction was quenched with 13 μL of 2X SDS-loading buffer. 5 μL of the final reaction was loaded onto a 4-20% polyacrylamide gel and was ran at 180 V for 35 minutes. The peptidoglycan product was transferred onto an Immune-Blot PVDF membrane (BioRad). The Lipid II product, labeled with BDL, was detected by incubation with streptavidin-IRdye (Li-COR1:10,000 dilution).

To quantify blots of biotinylated products from glycosyltransferase assays, lane profiles were plotted using the Fiji gel analyzer tool (Schindelin et al., 2012). Fragments larger than 48 kDa (the molecular weight of PBP4) were defined as long PG fragments. Fragments smaller than 48 kDa but larger than lipid II were defined as short PG fragments. The signal intensity from long PG fragments, short PG fragments, and lipid II were quantified and normalized to the total signal intensity in the lane. Results were analyzed using a two-way ANOVA, followed by Dunnett’s multiple comparisons test.

### Image Acquisition and Analysis

Conventional fluorescence and TIRF microscopy was performed on three distinct rigs. Rig #1 is described previously (Cho et al., 2016), and was used for the single-molecule tracking of MreB, as well as the TIRF:EPI determination. Rig #2 is a modified version of a previously described setup (Buss, Peters, Xiao, & Bernhardt, 2017), and was used for all non-TIRF fluorescence imaging, as well as the single-molecule tracking of sfGFP-PBP2 and sfGFP-PBP2(L61R). The new modifications include Ti-TIRF-EM Motorized Illuminator, a LUN-F laser launch with single fiber mode (488, 561, 640), Chroma TRF-EM 89901 Quad band set, Ti stage up kit, Sutter Emission filter wheel. Rig #3 was used for phase-contrast and DIC microscopy, and consists of a Nikon TE2000 microscope equipped with a 100x Plan Apo 1.4 NA objective, and a CoolSNAP HQ2 monochrome camera.

Sample preparation for all imaging was performed as described previously (Buss et al., 2017). Unless otherwise noted, cells were struck onto LB plates and inoculated in LB overnight prior to back-dilution (1:500) into M9 minimal media on the day of imaging. Induction of *Plac:^SW^mreB-mNeon* (attλHC897) was achieved with 100 μM IPTG throughout the duration of liquid growth. Imaging of *Plac:msfGFP-pbpa* (attHKHC943) and *Plac:msfGFP-pbpa(L61R)* (attHKPR128) required streaking onto M9 plates supplemented with 15 μM IPTG, followed by similar liquid growth. All conventional TIRF imaging was performed at 1s intervals for 1min duration with continuous illumination.

Analysis of phase-contrast images and conventional fluorescence was performed with Oufti and MATLAB (Paintdakhi et al., 2016). Single-molecule tracking data was analyzed with the Fiji plugin TrackMate, as described previously (Cho et al., 2016). We discarded single-molecule trajectories if they consisted of < 5 consecutive frames and had a minimum displacement of < 70 nm.

### SIM-TIRF Image Acquisition

We acquired SIM-TIRF images on the DeltaVision OMX SR (GE Healthcare Life Sciences). Imaging was performed at 37°C using ∼20 ms acquisitions (9 per frame, ∼200 ms total) at an interval of 1 s for 1-2 min duration. ^SW^MreB-mNeon polymer length was determined using custom MatLab software similar to that previously described (Buss, Coltharp, & Xiao, 2013). We only determined the lengths of polymers that were centrally positioned relative to the cell perimeter and believed to be entirely within the limited imaging area.

### TIRF:EPI Measurements

Epifluorescence illumination provides a wide depth-of-field (∼800 nm) and approximates the entire fluorescent population within a cell. TIRF illumination provides a narrow depth-of-field (∼200 nm) and approximates the membrane-associated population nearest the coverslip-sample interface. We assessed the relative abundance of the membrane-associated fraction of ^SW^MreB-mNeon within individual cells by calculating the ratio of the cumulative fluorescence intensity under TIRF and EPI. However, since TIRF intensity is highly affected by small changes in incident angle and z-focus, it is difficult to accurately compare separate TIRF:EPI datasets. Consequently we imaged both samples simultaneously. To differentiate the two strains, we expressed cytoplasmic mCherry (pAAY71) in either MG1655 attλHC897 or PR78 attλHC897 (Figure 6, supplement 1). We used data from both imaging pairs for analysis (Figure 6F).

### Strain Constructions

A complete list of strains can be found in Table 2.

**Table 2.** Strains used in this study

| Strain | Genotype<sup>a</sup> | Source/Reference<sup>b</sup> |
| --- | --- | --- |
| CAM333 | *E. coli* C43 ΔponB ΔpbpC ΔmtgA | Meeske et al., 2016 |
| dH5α(λpir) | *F*- *hsdR17 deoR recA1 endA1 phoA supE44 thi-1 gyrA96 relA1* $\Delta$(*lacZYA-argF*)U169 ϕ80d*lacZ*ΔM15 λpir | Laboratory stock |
| HC459 | MG1655 ΔpbpA::*kan* | Cho et al., 2014 |
| HC555 | MG1655 *yrdE*::*kan* | P1(λRed) x MG1655 |
| HC558 | MG1655 ΔpbpA*rodA*::*kan* | P1(λRed) x MG1655 |
| JAB576 | MG1655 *ybeM*::*frt mrdA*(L61R) *mreB*'-*mNeon*-'*mreB* Δ*yhdE*::*frt* | P1(HC583) (Cho et al., 2014) x PR78 |
| JAB593 | MG1655 *mreB*'-*mNeon*-'*mreB* | P1(HC583) (Cho et al., 2014) x MG1655 |
| MG1655 | *E. coli* *rph1 lvG rfb-50* | (Guyer et al., 1981) |
| MT4 | TB28 Δ*mreC*::*kan* | TB28 x P1(FB10)<br>(Bendezu et al., 2008) |
| PM7 | MG1655 Δ*ybeM2*::*kan* | Piet de Boer, unpublished |
| PM11 | MG1655 Δ*ybeM2*::*kan rodA*(A234T) | Piet de Boer, unpublished |
| PR5 | MG1655 *mreC*(R292H) *yrdE*::*kan* | P1(allelic exchange) x MG1655 |
| PR30 | MG1655 *mreC*(G156D) *yrdE*::*kan* | P1(allelic exchange) x MG1655 |
| PR55 | MG1655 Δ*ybeM1*::*kan* | P1(λRed) x MG1655 |
| PR68 | MG1655 Δ*ybeM1*::*kan pbpA*(L61R) | P1(allelic exchange) x MG1655 |
| PR78 | MG1655 Δ*ybeM1*::*frt pbpA*(L61R) | PR68/pCP20 |
| PR93 | MG1655 Δ*ybeM1*::*cat pbpA*(L61R) | P1(allelic exchange) x MG1655 |
| PR98 | MG1655 Δ*ybeM1*::*frt pbpA*(L61R) *yrdE*::*kan mreC*(G156D) | PR78 x P1(PR30) |
| PR99 | MG1655 Δ*ybeM1*::*frt pbpA*(L61R) *yrdE*::*kan mreC*(R292H) | PR78 x P1(PR5) |
| PR100 | MG1655 Δ*ybeM1*::*frt pbpA*(L61R) Δ*mreC*::*kan* | PR78 x P1(MT4) |
| PR101 | MG1655 Δ*ybeM1*::*cat* | P1(λRed) x MG1655 |
| PR115 | MG1655 Δ*ybeM1*::*cat pbpA*(T52A) | P1(suppressor strain) x MG1655 |
| PR116 | MG1655 ΔlysA::frt ΔpbpC::frt ΔmtgA::frt<br>ΔampD::frt mrcB(S247C) mrcA::frt ybeM1::cat | HC533 (Cho et al., 2016)<br>x P1(PR101) |
| PR117 | PR116 *pbpA*(L61R) | HC533 (Cho et al., 2016)<br>x P1(PR93) |
| PR124 | PR164 *pbpA*(T52A) *mreC*(R292H) | PR115 x P1(PR5) |
| PR125 | PR164 *pbpA*(T52A) *mreC*(G156D) | PR115 x P1(PR30) |
| PR127 | PR164 *pbpA*(L61R) | PR93 x P1(HC555) |
| PR128 | PR164 *pbpA*(L61R) *mreC*(R292H) | PR93 x P1(PR5) |
| PR129 | PR164 *pbpA*(L61R) *mreC*(G156D) | PR93 x P1(PR30) |
| PR131 | PR164 *pbpA*(T52A) | PR115 x P1(HC555) |
| PR132 | MG1655 ΔybeM1::frt | PR101/pCP20 |
| PR134 | MG1655 ΔrodZ::cat | P1(λRed) x MG1655 |
| PR136 | PR132 ΔmreBCD::kan | PR132 x P1(FB30)<br>(Bendezú et al., 2008) |
| PR137 | PR132 ΔmreCD::kan | PR132 x P1(FB14)<br>(Bendezú et al., 2008) |
| PR139 | PR132 *pbpA*(L61R) ΔmreBCD::kan | PR178 x P1(FB30)<br>(Bendezú et al., 2008) |
| PR140 | PR132 *pbpA*(L61R) ΔmreCD::kan | PR178 x P1(FB14)<br>(Bendezú et al., 2008) |
| PR142 | PR132 ΔrodZ::cat | PR132 x P1(PR134) |
| PR143 | PR143 *pbpA*(L61R) ΔrodZ::cat | PR78 x P1(PR134) |
| PR146 | MG1655 ΔlysA::frt ΔpbpC::frt ΔmtgA::frt<br>ΔampD::frt mrcB(S247C) mrcA::frt ybeM2::kan | HC533 (Cho et al., 2016)<br>x P1(PM7) |
| PR147 | PR146 *rodA*(A234T) | HC533 (Cho et al., 2016)<br>x P1(PM11) |
| PR149 | PR132 *pbpA*(L61R) ΔmreCD::kan ΔrodZ::cat | PR143 x P1(FB14)<br>(Bendezú et al., 2008) |
| PR150 | MG1655 ΔybeM2::frt | PM7/pCP20 |
| PR151 | PR150 *rodA*(A234T) | PM11/pCP20 |
| PR152 | PR150 ΔmreBCD::kan | PR150 x P1(FB30)<br>(Bendezú et al., 2008) |
| PR153 | PR150 $\Delta mreCD::kan$ | PR150 x P1(FB14)<br>(Bendezú et al., 2008) |
| PR154 | PR150 $\Delta rodZ::cat$ | PR150 x P1(PR142) |
| PR155 | PR150 $rodA(A234T) \Delta mreBCD::kan$ | PR151 x P1(FB30)<br>(Bendezú et al., 2008) |
| PR156 | PR150 $rodA(A234T) \Delta mreCD::kan$ | PR151 x P1(FB14)<br>(Bendezú et al., 2008) |
| PR157 | PR150 $rodA(A234T) \Delta rodZ::cat$ | PR151 x P1(PR142) |
| PR158 | MG1655 $\Delta ybeM2::frt yrdE-kan$ | PR150 x P1(HC555) |
| PR159 | PR158 $mreC(R292H)$ | PR150 x P1(PR5) |
| PR160 | PR158 $mreC(G156D)$ | PR150 x P1(PR30) |
| PR161 | PR158 $rodA(A234T)$ | PR151 x P1(HC555) |
| PR162 | PR158 $rodA(A234T) mreC(R292H)$ | PR151 x P1(PR5) |
| PR163 | PR158 $rodA(A234T) mreC(G156D)$ | PR151 x P1(PR30) |
| PR164 | MG1655 $\Delta ybeM1::cat yrdE-kan$ | PR101 x P1(HC555) |
| PR165 | PR164 $mreC(R292H)$ | PR101 x P1(PR5) |
| PR166 | PR164 $mreC(G156D)$ | PR101 x P1(PR30) |
| Lemo21(λDE3) | $fhuA2 [lon] ompT gal (\lambda DE3) [dcm] \Delta hsdS/ pLemo$ | Wagner et al., 2008 |
| SM10(λpir) | $KanR thi-1 thr leu tonA lacY supE recA::RP4-2-Tc::Mu att\lambda::pir$ | Miller & Mekalanos, 1988 |
| TB10 | MG1655 $\lambda \Delta cro-bio nad::Tn10$ | Johnson, Lackner, Hale,<br>& de Boer, 2004 |
| TB28 | MG1655 $\Delta lacIZYA::frt$ | Bernhardt & de Boer,<br>2004 |

<sup>a</sup> The Kan<sup>R</sup> cassette is flanked by *frt* sites for removal by FLP recombinase. An *frt* scar remains following removal of the cassette using FLP recombinase expressed from pCP20.
<sup>b</sup> Strain constructions by P1 transduction are described using the shorthand: P1(donor) x recipient. Transductants were selected on LB Kan, LB Tet, or LB Cm plates where appropriate. $\lambda$ Red indicates strains constructed by recombineering (see Experimental Procedures for details). Strains resulting from the removal of a drug resistance cassette using pCP20 are indicated as: Parental strain/pCP20.

HC555 [MG1655 *yrdE-kan*]-A Kan^R^ cassette was inserted in the intergenic space downstream of *yrdE* (genotype designated *yrdE-kan* in this paper), so that it could be used to co-transduce the *mre* locus. The KanR cassette was amplified from pKD13 (Datsenko & Wanner, 2000) using primers o1141 (<u>TGGCGCTAATTTCGTGAATTGTGCGGCTTGTTGCAAATTA</u>ATTCCGGGGATCCGTCGACC) and o1142 (<u>ATAATCAACAGCTAACATGTAAATAACCTTCAACACCGT</u>GTGTAGGCTGGAGCTGCTTCG). The resulting PCR product was purified and electroporated into recombineering strain TB10 (using the same protocol as described for recombineering with DY330 (Yu et al., 2000)), and recombinants were selected at 30℃ on LB agar supplemented with 25 μg/mL kanamycin. The *yrdE-kan* allele was moved from this strain into MG1655 by P1-mediated transduction, generating strain HC555. The growth rate and cell dimensions of this strain are indistinguishable from wild type.

PR5 [MG1655 *mreC(R292H) yrdE-kan*]*-*A strain harboring the chromosomal *mreC(R292H)* mutation was constructed by allelic exchange, using a previously described protocol (Philippe, Alcaraz, Coursange, Geiselmann, & Schneider, 2004). The *pir-*dependent suicide plasmid pPR84 [*sacB Cm^R^*] was introduced into the recipient strain HC555/pTB63 [*yrdE-kan Tet^R^*] by conjugative transfer from the donor strain SM10(λpir). Briefly, 5 mL of exponential-phase cultures (OD_600_ ≈ 0.3) of the donor and recipient strains were filtered onto the same 0.2 μm PES filter. This filter was placed cell-side-up on an LB agar plate and incubated for four hours at 37℃. Cells from the filter were then resuspended in 1 mL of LB, then plated on LB agar supplemented with chloramphenicol and tetracycline, and incubated at 30℃ for 24 hours to select for exconjugants that contain pPR84 integrated into the chromosome via a single cross-over. Exconjugants were streaked on the same medium, and screened to identify isolates with spherical cell shape (indicating that the cross-over had occurred at the *mre* locus, resulting in *mre(R292H)* expression). An exconjugant colony was resuspended in LB, serially diluted, plated on LB agar lacking NaCl and supplemented with 6% sucrose, and incubated at 30℃ for 24 hours to select for recombinants that have lost the *sacB-*containing plasmid via a single cross-over. Sucrose-resistant colonies were replica-streaked on LB agar with and without chloramphenicol. Sucrose-resistant, chloramphenicol-sensitive isolates were screened for spherical cell morphology, indicating that *mreC(R292H)* had replaced the wild-type allele of *mreC* at the native chromosomal locus. This was confirmed by PCR followed by Sanger sequencing. Strain PR5 was obtained by P1-mediated transduction of the genomic region near *yrdE-kan* (including *mre(R292H)*) from the primary isolate into MG1655. Transductants were selected on M9 agar supplemented with casamino acids, glucose, and kanamycin, screened for spherical cell shape, and confirmed by PCR and sequencing of *mreC*.

PR30 [MG1655 *mreC(G156D) yrdE-kan*] was constructed by allelic exchange using the suicide vector pPR93, following the same protocol as described above for PR5.

PR55 [MG1655 *ΔybeM1::kan*]-A Kan^R^ cassette was used to replace the *ybeM* pseudogene, so that this marker could be used to co-transduce the *mrd* locus. The Kan^R^ cassette was amplified from pKD13 (Datsenko & Wanner, 2000) using primers o1237 (<u>TCGTTGGCGAATTTTACGACTCTGACAGGAGGTGGCAATG</u>ATTCCGGGGATCCGTCGACC) and o1238 (<u>AGCGCCGAGTAAAAAAACATCATAATAATTGCGGCGGCGC</u>GTGTAGGCTGGAGCTGCTTCG). The resulting PCR product was purified and electroporated into recombineering strain TB10 (using the same protocol as described for recombineering with DY330 (Yu et al., 2000)), and recombinants were selected at 30℃ on LB agar supplemented with 25 μg/mL kanamycin. The *ΔybeM::kan* allele was moved from this strain into MG1655 by P1-mediated transduction, generating strain PR55. The growth rate and cell dimensions of this strain are indistinguishable from wild type.

PR68 [MG1655 *ΔybeM1::kan pbpA(L61R)*]*-*A strain harboring the chromosomal *pbpA(L61R*) mutation was constructed by allelic exchange, using a previously described protocol (Philippe et al., 2004). The *pir-*dependent suicide plasmid pPR101 [*sacB Cm^R^*] was introduced into the recipient strain PR55/pTB63 [*ΔybeM1::kan / Pnative::ftsQAZ Tet^R^*] by conjugative transfer from the donor strain SM10(λpir). Exconjugants that had integrated the plasmid into the genome via a single cross-over were selected on medium containing chloramphenicol and tetracycline. Exconjugants were then plated on sucrose to select for loss of the plasmid via a second recombination event. Suc^R^ Cm^S^ colonies were screened by PCR and sequencing for the presence of the *pbpA(L61R)* mutation. Strain PR68 was obtained by P1-mediated transduction of the genomic region near *ΔybeM::kan* (including *pbpA(L61R)*) from the primary isolate into MG1655.

PR101 [MG1655 *ΔybeM1::cat*]*-*A Cm^R^ cassette was used to replace the *ybeM* pseudogene, so that this marker could be used to co-transduce the *mrd* locus. The Cm^R^ cassette was amplified from pKD3 (Datsenko & Wanner, 2000) using primers o1415 (<u>TCGTTGGCGAATTTTACGACTCTGACAGGAGGTGGCAATG</u>CATATGAATATCCTCCTTAG) and o1416 (<u>AGCGCCGAGTAAAAAAACATCATAATAATTGCGGCGGCGC</u>GTGTAGGCTGGAGCTGCTTC). These primers are designed such that the *ΔybeM1::cat* lesion is identical to the *ΔybeM1::kan* lesion in PR55, the only difference being the antibiotic resistance cassette. The resulting PCR product was purified and electroporated into recombineering strain TB10 (using the same protocol as described for recombineering with DY330 (Yu et al., 2000)), and recombinants were selected at 30℃ on LB agar supplemented with 25 μg/mL chloramphenicol. The *ΔybeM::cat* allele was moved from this strain into MG1655 by P1-mediated transduction, generating strain PR101. The growth rate and cell dimensions of this strain are indistinguishable from wild type.

PR93 [MG1655 *ΔybeM1::cat pbpA(L61R)*]*-*The *ΔybeM1::cat* cassette was transferred from donor strain PR101 to recipient strain PR68 [MG1655 *ΔybeM1::kan pbpA(L61R)*] by P1-mediated transduction. Since *ybeM* and *pbpA* are closely linked, most transductants contained the wild-type *pbpA* sequence from donor strain PR101. PCR and sequencing were used to identify a rare Kan^S^ Cm^R^ transductant that retained the *pbpA(L61R)* sequence.

PR115 [MG1655 *ΔybeM1::cat pbpA(T52A)*]*-*This strain was constructed in a two-step procedure. First, the *ΔybeM1::cat* cassette from PR101 was transduced into a suppressor strain derived from PR30 [*mreC(G156D)*] that contains the spontaneous mutation *pbpA(T52A).* Although *ybeM* and *pbpA* are closely linked, all transductants retained the *pbpA(T52A)* mutation, because this mutation permits survival on LB. P1 lysates were prepared on this intermediate strain, and the *ΔybeM1::cat pbpA(T52A)* locus was co-transduced into MG1655, generating strain PR115. The presence of the *pbpA(T52A)* mutation was confirmed by PCR and sequencing.

PM7 [MG1655 Δ*ybeM2::kan*] was a gift from Dr. Piet de Boer. This strain contains a kanamycin resistance cassette in the *ybeM* locus. Since the exact junction points are different from those in PR55 [Δ*ybeM1::kan*], the allele is designated Δ*ybeM2::kan*.

PM11 [MG1655 Δ*ybeM2::kan rodA(A234T)*] contains a *rodA(A234T)* mutation in the PM7 genetic background. This strain was a gift from Dr. Piet de Boer.

PR134 [MG1655 Δ*rodZ::cat*]-A Cm^R^ cassette was used to replace the region between the 2nd codon and 7th codon from the stop codon of *rodZ,* as described previously (Baba et al., 2006; Yu et al., 2000). The Cm^R^ cassette was amplified from pKD3 (Datsenko & Wanner, 2000) using primers o1953 (<u>CTCCCGCGTTACCCGTCTGTTACTGCGCCGGTGATTGTTC</u>GTGTAGGCTGGAGCTGCTTC) and o1954 (<u>CGGCATCTCAATTCTCATTTAAACGTACCTGCAGCGAATG</u>CATATGAATATCCTCCTTAG). The resulting PCR product was purified and electroporated into MG1655/pKD46 as described previously (Datsenko & Wanner, 2000), and recombinants were selected at 42℃ on M9 agar supplemented with casamino acids, glucose, and 25 μg/mL chloramphenicol. PR134 was made by P1 transduction of Δ*rodZ::cat* from this intermediate strain into an MG1655 recipient strain.

HC558 [MG1655 Δ*pbpArodA::kan*]-A Kan^R^ cassette was used to replace the region between the 2nd codon of *pbpA* and 5th codon from the stop codon of *rodA,* as described previously (Yu et al., 2000). The Kan^R^ cassette was amplified from pKD13 (Datsenko & Wanner, 2000) using primers o1094 (<u>TGAGTGATAAGGGAGCTTTGAGTAGAAAACGCAGCGGATG</u>ATTCCGGGGATCCGTCGACC) and o1095 (<u>CCACTGCTTACGCATTGCGCACCTCTTACACGCTTTTCGA</u>TGTAGGCTGGAGCTGCTTCG). The resulting PCR product was purified and electroporated into TB10/pCX16, using the same protocol as described for recombineering with DY330 (Yu et al., 2000)), and recombinants were selected at 30℃ on LB agar supplemented with 25 μg/mL kanamycin. HC558/pRY47, HC558/pHC857, and HC558/pSS43 were made by P1-mediated transduction of Δ*pbpArodA::kan* from this intermediate strain into MG1655 containing the corresponding plasmid.

### Plasmid constructions

A complete list of plasmids can be found in Table 3.

**Table 3.**
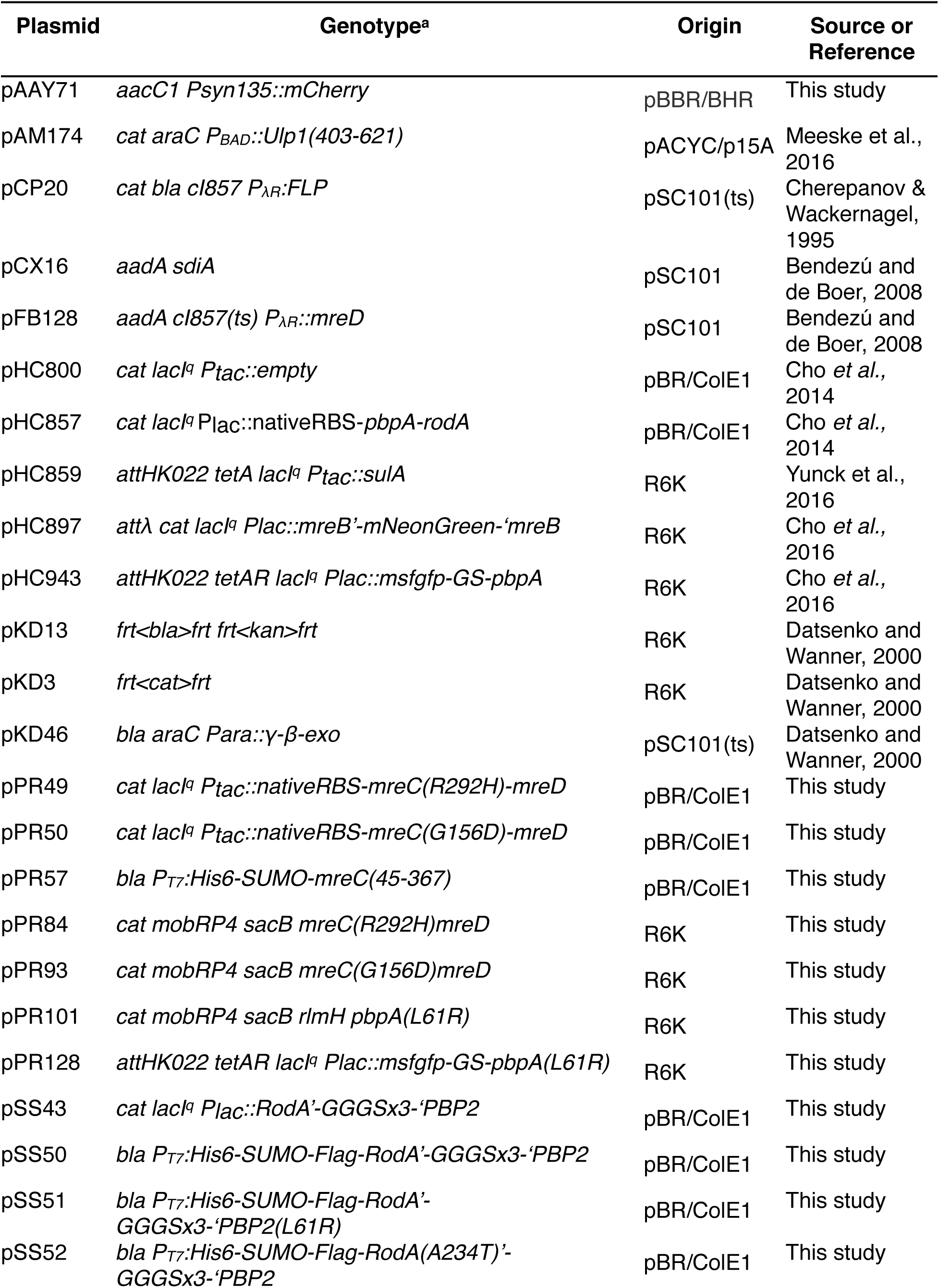
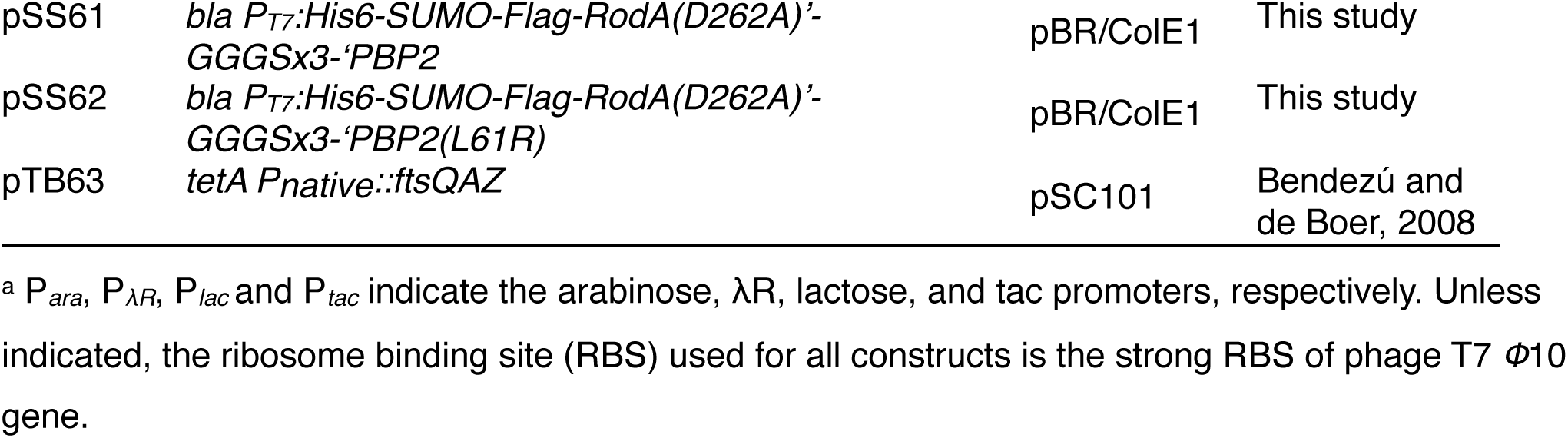
Plasmids used in this study

pPR49 [*colE1 cat lacI^q^ Ptac::nativeRBS-mreC(R292H)-mreD*]*-*Primers o882 (GTCA*<u>TCTAGA</u>*CTGCCTGGTCTGATACGAGAATACGCATAACTTATG), o918 (CTGCATCAG**ATG**TTCATTAGCAACACGATGC), o919 (GCTAATGAA**CAT**CTGATGCAGATGATGCCGC), and o905 (GTCA*<u>AAGCTT</u>*TTATTGCACTGCAAACTGCTGACGG) were used to amplify MG1655 genomic DNA and introduce the R292H mutation into *mreC* using overlap-extension PCR. The product was PCR purified, digested with XbaI/HindIII, and cloned into similarly digested pHC800.

pPR50 [*colE1 cat lacI^q^ Ptac::nativeRBS-mreC(G156D)-mreD*]*-*Primers o882, o914 (GACCAACAAC**ATC**TTTGTCGCTGATGACCGGC), o915 (GCGACAAA**GAT**GTTGTTGGTCAGGTGGTGG), and o905 were used to amplify MG1655 genomic DNA and introduce the G156D mutation into *mreC* using overlap-extension PCR. The product was PCR purified, digested with XbaI/HindIII, and cloned into similarly digested pHC800.

pPR57 [*colE1 bla P_T7_:His6-SUMO-mreC(45-367)*]*-*Primers o883 (GTCA*<u>AAGCTT</u>*CTATTGCCCTCCCGGCGCAC) and o920 (ATTGGT*<u>GGATCC</u>*GCCGTCAGTCCTTTCTACTTTGTTTCC) were used to amplify the insert (BamHI-mreC(45-367)-HindIII) from MG1655 genomic DNA. This insert was cut with BamHI/HindIII and ligated into similarly digested pTD68 (Uehara, Parzych, Dinh, & Bernhardt, 2010).

pPR84 [*cat mobRP4 sacB mreC(R292H)mreD*]*-*Primers o1157 (GTCA*<u>GAGCTC</u>*CTGCCTGGTCTGATACGAG) and o1158 (GTCA*<u>TCTAGA</u>*TTATTGCACTGCAAACTGCTGACGG) were used to amplify the insert (SacI-mreC(R292H)-mreD-XbaI) from pPR49. This insert was cut with SacI/XbaI and ligated into similarly digested pDS132 (Philippe et al., 2004).

pPR93 [*cat mobRP4 sacB mreC(G156D)mreD*]*-*Primers o1157 and o1158 were used to amplify the insert (SacI-mreC(G156D)-mreD-XbaI) from pPR50. This insert was cut with SacI/XbaI and ligated into similarly digested pDS132 (Philippe et al., 2004).

pPR101 [*cat mobRP4 sacB rlmH pbpA(L61R)*]*-*Primers o1285 (GTCA*<u>GAGCTC</u>*CATCCGCTGGTTCGCGTGCTGG) and o1286 (GTCA*<u>TCTAGA</u>*TCCCCATATCGTAGGCCACCTG) were used to amplify the insert (a segment of genomic DNA encompassing a 3’ fragment of *rlmH* and the 5’ half of *pbpA(L61R),* flanked by SacI and XbaI restriction sites) from a suppressor mutant derived from PR5, containing the spontaneous mutation *pbpA(L61R)*. This insert was cut with SacI/XbaI and ligated into similarly digested pDS132 (Philippe et al., 2004).

pPR128 [*attHK022 tetAR lacIq Plac::msfgfp-GS-pbpA(L61R)*]-*pbpA(L61R)* was PCR amplified from PR68 gDNA using primers o264 (GCTAAAGCTTTTTATTCGGATTATCCGTCATG) and o1041 (GCTA*<u>GGATCC</u>*AAACTACAGAACTCTTTTCGCGACTATACG). The resulting PCR product was digested with BamHI and HindIII restriction enzymes and cloned into pHC943, which was pre-digested with the same enzymes.

pSS43 [*colE1 cat lacI^q^ Plac::RodA’-GGGSx3-’PBP2*] was generated in two steps. First, the insert containing RodA was amplified from MG1655 genomic DNA as a template with primers oSS37 (TCGACAAGCTTTTACACGCTTTTCGACAACATTTTCCTGTGG) and oSS59 (GTTTAACTTTAAGAAGGAGATATACCATGACGGATAATCCGAATAAAAAAACATTCTGGG). The resulting PCR product was then assembled with XbaI/HindIII-digested pRY47 [*colE1 cat lacI^q^ Plac::empty*] using the isothermal assembly procedure (Gibson et al., 2009). This intermediate plasmid was amplified using primers oSS62 (CCGCAGCGGAGGACCATTAAGCTTGTCACCGATACGCGAGCGAACGTGAAGCGACTGCTG) and oSS75 (AGAACCGCCACCGGAGCCACCGCCGCTACCGCCACCCACGCTTTTCGACAACATTTTCCT) to create the vector for isothermal assembly with an insert containing GGGSx3-’PBP2, amplified from MG1655 genomic DNA with primers oSS61 (CTCGCGTATCGGTGACAAGCTTAATGGTCCTCCGCTGCGGCAACCGCTGGATTTTCCGCA) and oSS74 (GGTGGCGGTAGCGGCGGTGGCTCCGGTGGCGGTTCTAAACTACAGAACTCTTTTCGCGAC).

pSS50 [*colE1 bla P_T7_:His6-SUMO-Flag-RodA’-GGGSx3-’PBP2*] was generated in a two-piece isothermal assembly reaction with an insert containing RodA’-GGGSx3-’PBP2, which was amplified from pSS43 [*colE1 cat lacIq Plac::RodA’-GGGSx3-’PBP2*] with oligonucleotide primers oSS82 (GGGTCATCCACGGATAATCCGAATAAAAAAACATTCTGGGATAAAGTCCATCTCGATCCC) and oSS84 (GCAGCCGGATCCCCTTCCTGCAGTCACCCGGGCTTAATGGTCCTCCGCTGCGGCAACCGC), and pAM172 [*colE1 bla P_T7_::His6-SUMO-Flag-RodA*] (Meeske et al., 2016), which was amplified with oligonucleotide primers oSS83 (TCCCAGAATGTTTTTTTATTCGGATTATCCGTGGATGACCCCCCAGGGCCTTGAAACAAC) and oSS85 (AATCCAGCGGTTGCCGCAGCGGAGGACCATTAAGCCCGGGTGACTGCAGGAAGGGGATCC).

pSS51 [*colE1 bla P_T7_:His6-SUMO-Flag-RodA’-GGGSx3-’PBP2(L61R)*] was generated in a two-piece isothermal assembly reaction with an insert containing PBP2(L61R), which was amplified from PR68 gDNA with oligonucleotide primers oSS74 and oSS84, and pSS50, which was amplified with oligonucleotide primers oSS75 and oSS85.

pSS52 [*colE1 bla P_T7_:His6-SUMO-Flag-RodA(A234T)’-GGGSx3-’PBP2*] was generated in a two-piece isothermal assembly reaction with an insert containing RodA(A234T), which was amplified from PR151 with oligonucleotide primers oSS75 and oSS82, and pSS50, which was amplified with oligonucleotide primers oSS74 and oSS83.

pSS60 [*colE1 bla PT7:His6-SUMO-Flag-RodA(D262A)’-GGGSx3-’PBP2*] was generated in a two-piece isothermal assembly reaction with an insert containing RodA(D262A), which was generated by overlap extension PCR using oligonucleotide primers oSS36 (TTGGTGGATCCATGACGGATAATCCGAATAAAAAAACATTCTGGG), oSS37, oSS96 (ACGCCATACTGCCTTTATCTTCGCGGTACTGGC), and oSS97 (CGAAGATAAAGGCAGTATGGCGTTCGGGGAGAA), and pSS50, which was amplified with oligonucleotide primers oSS74 and oSS83.

pSS62 [*colE1 bla PT7:His6-SUMO-Flag-RodA(D262A)’-GGGSx3-’PBP2(L61R)*] was generated in a two-piece isothermal assembly reaction with an insert containing RodA(D262A), which was generated by overlapping PCR using oligonucleotide primers oSS36, oSS37, oSS96, and oSS97, and pSS51, which was amplified with oligonucleotide primers oSS74 and oSS83.

pAAY71 [*aacC1 Psyn135::mCherry*]-To a vector for expressing cytoplasmic mCherry, the mCherry gene was PCR-amplified from pAAY65 [*aacC1 Psyn135::ssdsbA-mCherry*] (Yakhnina, McManus, & Bernhardt, 2015) template using primers oAAY1 (TTTTCATATGTCCAAGGGCGAGGAGGATAACCTG) and oAAY2 (TTTTGTCGACTTATTAGGATCCGCCAGCACCTTTGTAC). The resulting PCR product was digested with NdeI and SalI restriction enzymes and cloned into pAAY65, which was pre-digested with the same enzymes.

## ACKNOWLEDGEMENTS

The authors would like to thank all members of the Bernhardt, Rudner, Walker, Kahne, Kruse, and Garner labs for advice and helpful discussions. Piet de Boer kindly provided us with the *rodA(A234T)* allele. This work was supported by the National Institute of Allergy and Infectious Diseases of the National Institutes of Health (R01AI099144 to TGB and SW, and CETR U19 AI109764 to TGB, DK, SW, and ACK), and the Canadian Institutes of Health Research (Doctoral Research Award to PDAR).

---

**Figure 3, Supplement 1:**
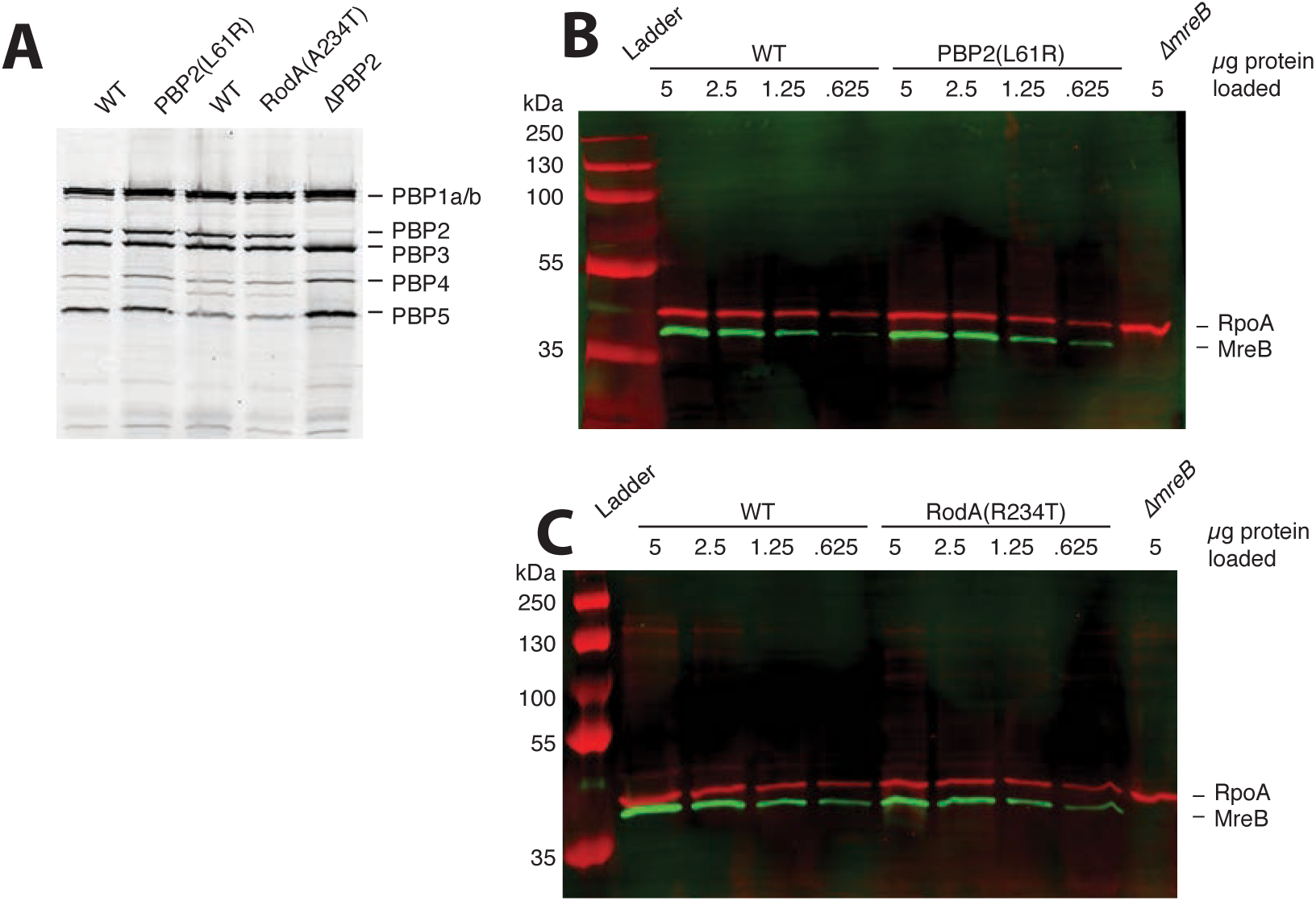
MreB and PBP2 levels are unaffected in the *pbpA\** mutant. **A.** Overnight cultures of each strain [PR132, PR78, PR150, PR151, TU230/pTB63] were diluted 1/200 and grown until the OD_600_=0.3, then labelled with Bocillin as described in methods. Membrane fractions were isolated, and 15 μg of total protein was loaded in each lane of a 10% SDS-PAGE gel. **B.** Western blot detecting RpoA (red) and MreB (green). Each lane contains the indicated amount of total protein from exponential-phase (OD_600_=0.3) whole cell extracts of WT [PR132], *pbpA(L61R)* [PR78], and Δ*mreBCD::kan* [TU233/pTB63]. **C.** Western blot detecting RpoA (red) and MreB (green). Each lane contains the indicated amount of total protein from exponential-phase (OD_600_=0.3) whole cell extracts of WT [PR150], *rodA(A234T)* [PR151], and Δ*mreBCD::kan* [TU233/pTB63].

**Figure 3, supplement 2:**
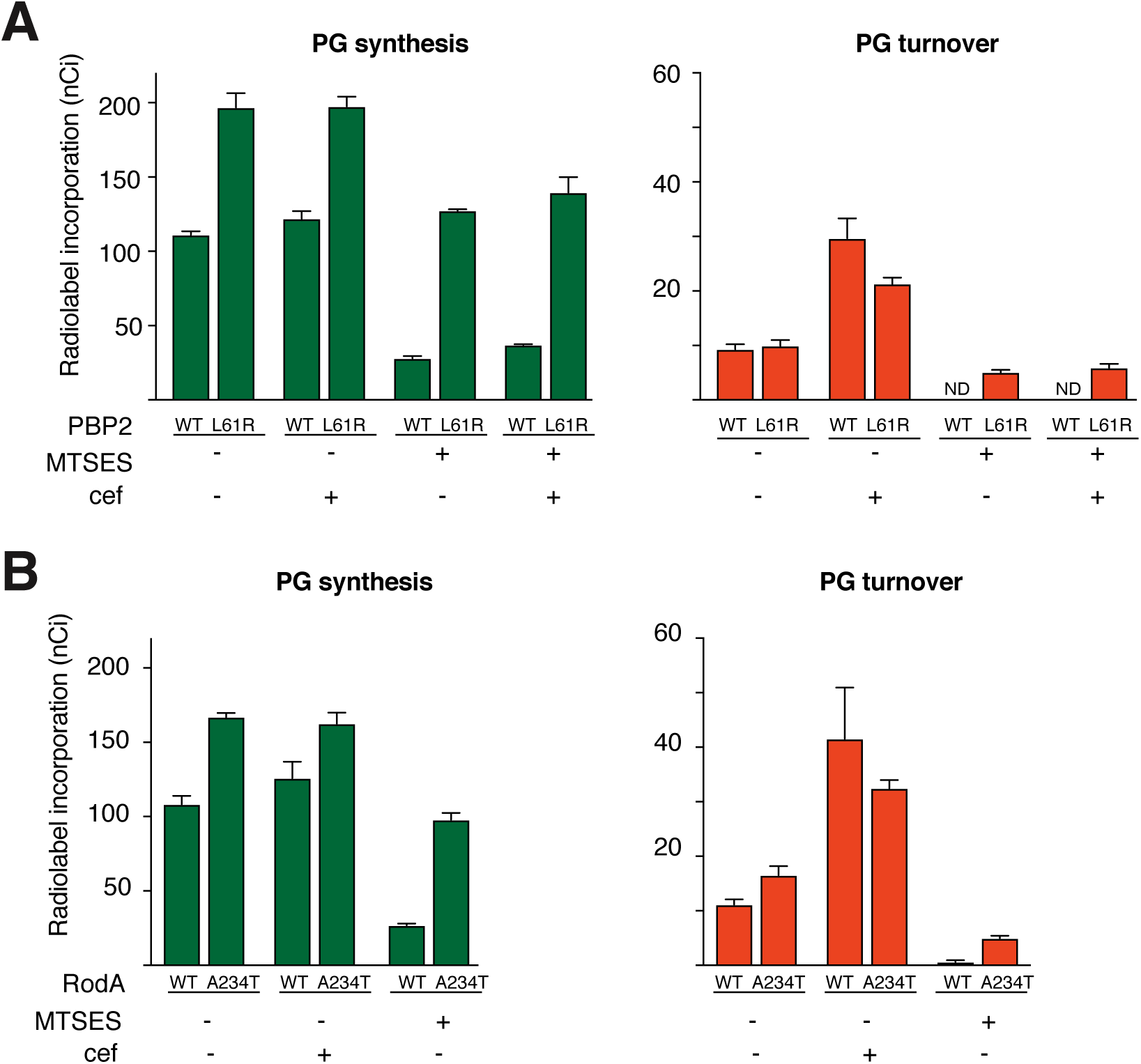
Increased synthesis in *pbpA rodA* mutants is independent of aPBP activity. **A.** Labeling strains encoding PBP2(WT) or PBP2* at the native genomic locus [PR116(attHKHC859) and PR117(attHKHC859)] were pre-treated with 1.5 mM IPTG to induce SulA production, and 1 mM MTSES and/or 100 μg/mL cefsulodin, as indicated. Strains were then pulse-labelled with [3H]-mDAP, and peptidoglycan precursors (UDP-MurNAC-pentapeptide), synthesis, and turnover products (anhydroMurNAC-tripeptide and -pentapeptide) were measured. Results are the average of three independent experiments. Error bars represent the standard error of the mean. **B.** The same experiments and analysis as in (A) were performed using labeling strains encoding RodA(WT) or RodA* at the native genomic locus [PR146(attHKHC859) and PR147(attHKHC859)].

**Figure 5, supplement 1:**
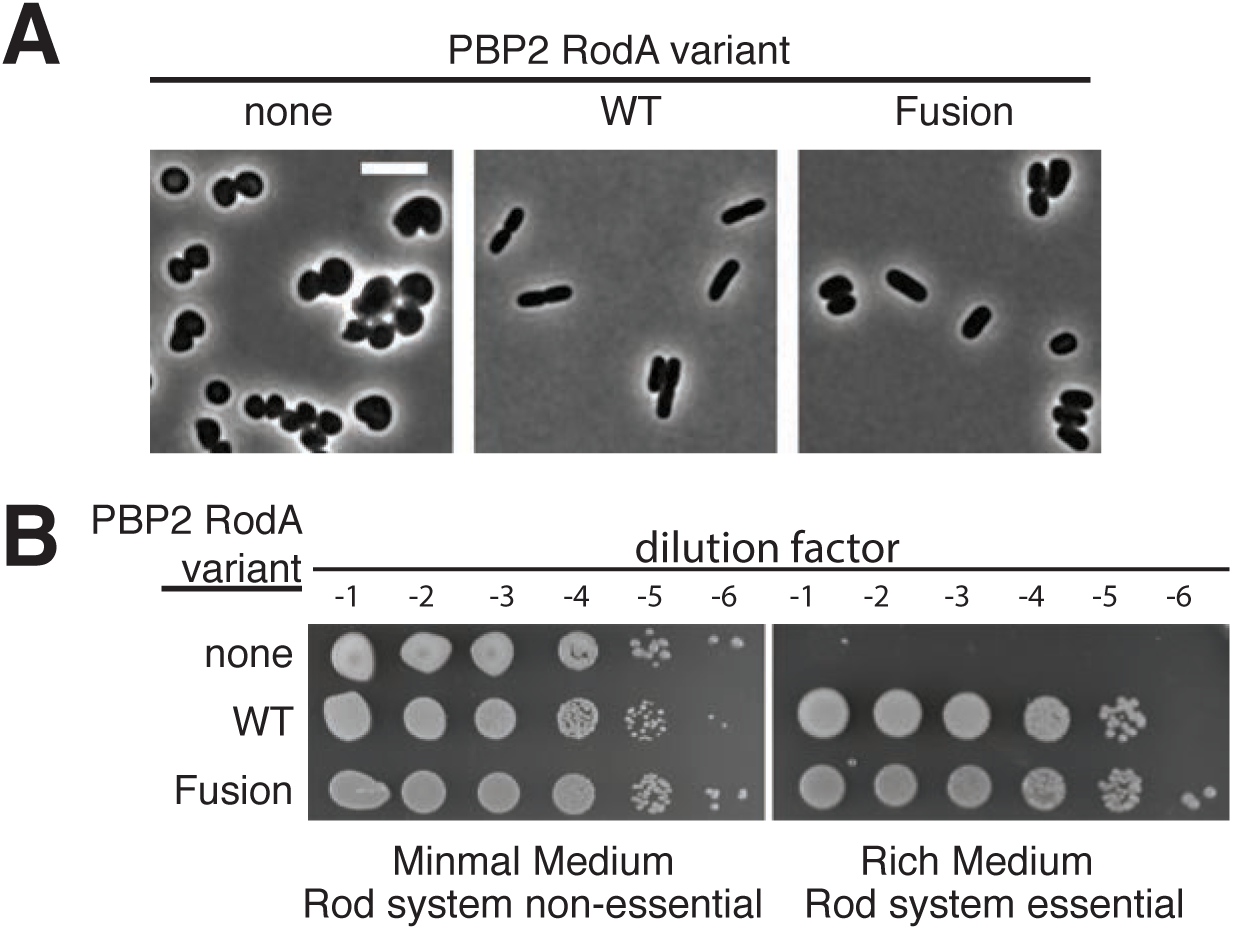
RodA-PBP2 fusion complements *ΔpbpA rodA* shape defect. A. Overnight cultures of *ΔpbpArodA::Kan* cells [HC558] harboring vectors expressing the indicated genes under P_lac_ control [pRY47, pHC857, pSS43] were diluted to OD_600_=0.005 in 3 mL of M9 medium supplemented with 0.2% casamino acids, 0.2% maltose, and 25 μM IPTG. When the OD_600_ reached 0.1-0.2, cells were fixed and imaged using phase contrast microscopy. Scale bar, 5 μm. B. Overnight cultures of the above strains were serially diluted and spotted on either M9 agar supplemented with 0.2% casamino acids and 0.2% maltose, or LB agar containing 50 μM IPTG.

**Figure 5, supplement 2:**
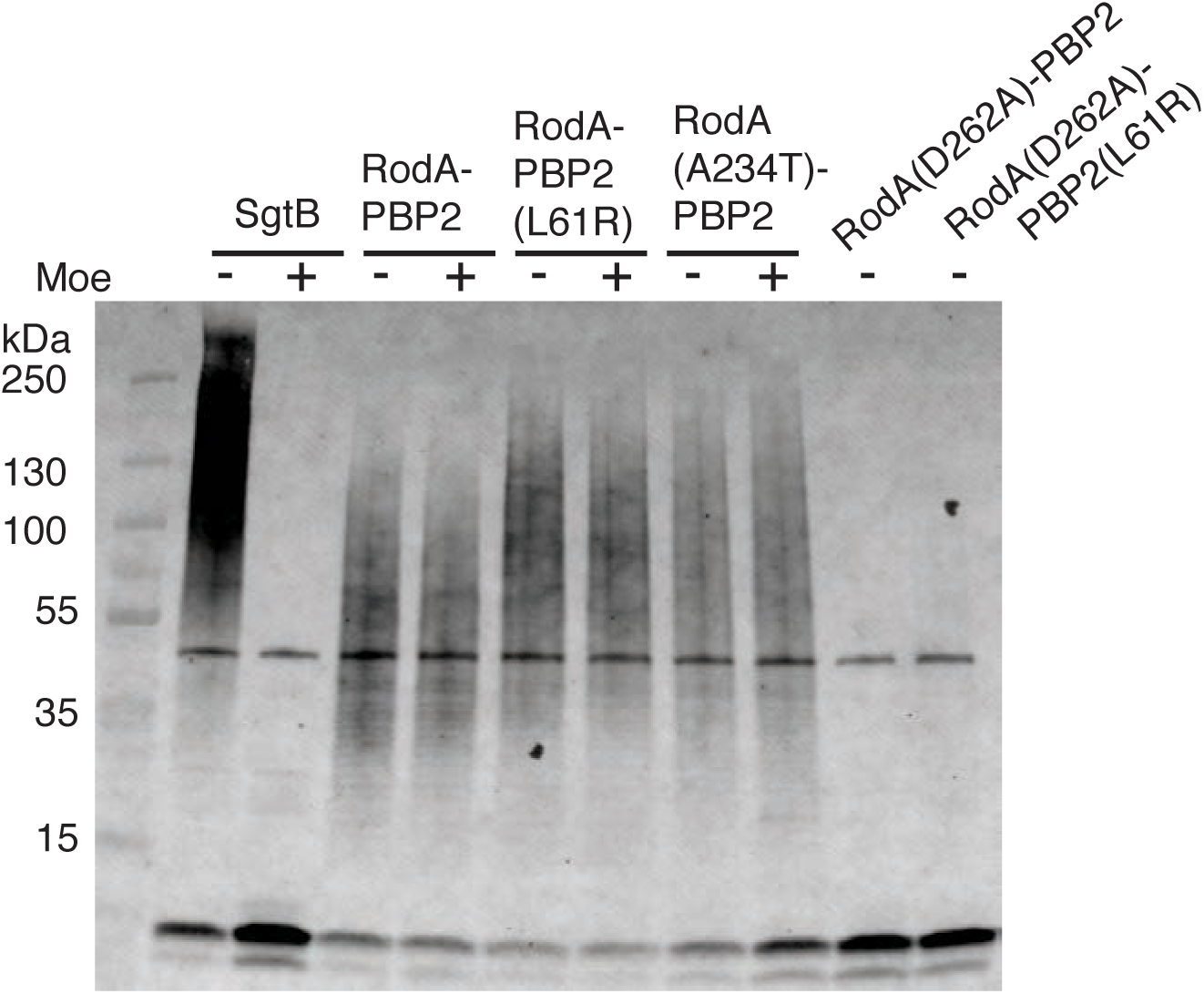
RodA is the only glycosyltransferase present in the reactions. Blot detecting the peptidoglycan products produced by the RodA-PBP2 fusion constructs from the glycosyltransferase assays using *E. coli* Lipid II. The product was detected by BDL labeling with *S. aureus* PBP4. Glycosyltransferase activity was assessed in the presence and absence of moenomycin (moe). All reactions were analyzed after 20 min. SgtB, a moenomycin-sensitive glycosyltransferase purified from *S. aureus*, was used as a positive control. The introduction of a point mutation into RodA(D262A) disrupts the production of the polymerization product.

**Figure 6, supplement 1:**
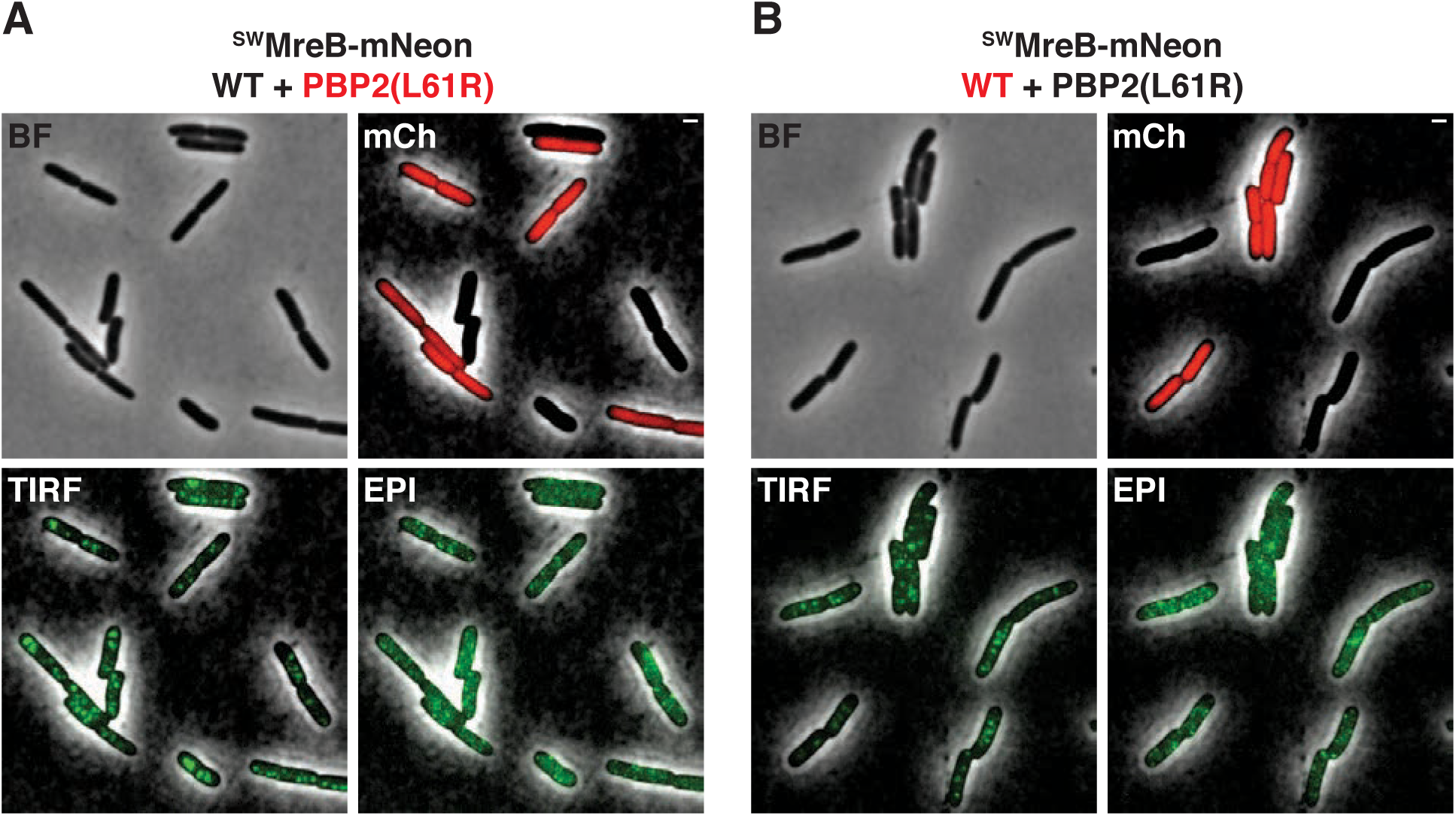
TIRF:EPI measurements. **A.** Representative micrographs of a mixed population containing MG1655(attλHC897) and PR78(attλHC897)/pAAY71(P_syn135_:mCherry). Images are presented as phase-contrast (BF) and fluorescence images overlaid with a contrast-adjusted phase-contrast image. Cytoplasmic mCherry (mCh) is pseudocolored red, while MreB-^SW^mNeon is pseudocolored green and labeled according to its illumination setting (TIRF, EPI). Scale bars, 1μm. **B.** Same as above, but with the mixed populations containing PR78(attλHC897) and MG1655(attλHC897)/pAAY71(P_syn135_:mCherry).

Supp. Movie #1 - Conventional TIRF microscopy of MreB-^SW^mNeon in MG1655

Supp. Movie #2 - Conventional TIRF microscopy of MreB-^SW^mNeon in PR78

Supp. Movie #3 - Conventional TIRF microscopy of msfGFP-PBP2 in MG1655

Supp. Movie #4 - Conventional TIRF microscopy of msfGFP-PBP2(L61R) in MG1655

Supp. Movie #5 - SIM-TIRF microscopy of MreB-^SW^mNeon in MG1655

Supp. Movie #6 - SIM-TIRF microscopy of MreB-^SW^mNeon in PR78

